# Integration of artificial intelligence and high-content screening enabled identification of drugs for long-term treatment of cerebral cavernous malformation disease

**DOI:** 10.64898/2025.12.08.693036

**Authors:** Eduardo Frias-Anaya, Helios Gallego-Gutierrez, Cassandra Bui, Janeth Ochoa Birrueta, Jeffrey Steinberg, Ingrid Reynolds Niesman, Brendan Gongol, Brina Nguyen, Aaryaman Sawhney, Zaida Mizushima, Nyle Connolly, Bethan Kilpatrick, Issam A. Awad, Hemal H. Patel, JoAnn Trejo, Jeremiah D. Momper, Andrea Taddei, Miguel Alejandro Lopez-Ramirez

**Author notes:** Correspondence should be addressed M.A.L.R, 9500 Gilman Diver, BSB 5096, La Jolla, CA 92093. Contributed equally (Eduardo Frias-Anaya, Helios Gallego-Gutierrez, Cassandra Bui).

## Abstract

**Background:** Adults and children with cerebral cavernous malformations (CCMs) are at risk of experiencing lifelong complications such as hemorrhagic strokes, neurological deficits, and epileptic seizures. These complications can severely reduce quality of life. At present, there is no safe or effective therapeutic option for the long-term treatment of CCMs.

**Methods:** Using advanced artificial intelligence (AI) and machine learning models, powered by the Benevolent Platform™, we aimed to identify therapeutic drug targets for CCM pathology (e.g., CCM1, CCM2, CCM3). An AI integrative approach utilized various data types from biomedical entities, including diseases, genes, tissues, and biological mechanisms, together with CCM transcriptomic experimental data. High-throughput drug screening of AI-selected FDA-approved medications, analyses of mitochondrial morphology, and studies on pharmacokinetics, pharmacodynamics, and toxicology were conducted in CCM animal models to identify drugs that could potentially be repurposed for the long-term treatment of CCM disease.

**Results:** AI predicted the AMPK (AMP-activated protein kinase) and mTOR (mammalian target of rapamycin) pathways as potential therapeutic targets that contribute to CCM pathology. High-content screening validation revealed that the FDA-approved drug metformin, which acts as an AMPK agonist and mTOR inhibitor, can reverse changes in cell-cell junction organization and increase KLF4 expression, a marker for CCM, in human CCM endothelial cells in cultured assays. In addition, pharmacodynamic markers of metformin were observed in CCM mouse models (*Slco1c1-iCreERT2;Krit1^fl/fl^;Pten^fl/wt^* and *Slco1c1-iCreERT2;Pdcd10^fl/fl^*) including reduced S6 kinase or ribosomal protein phosphorylation, a marker of decrease mTOR signaling, and increased AMPK phosphorylation, a marker of AMPK activation, that corresponded to reduced lesion burden. Pharmacokinetic and toxicological studies in CCM animal models showed that that metformin penetrates the brain and long-term administration has a favorable safety profile. We also demonstrated that brain endothelial cells in chronic CCM mouse models exhibit increased levels of the inflammatory marker VCAM-1, which is associated with altered mitochondrial phenotypes, as observed by immunofluorescence, MITO-tagging, and electron microscopy analysis. Additionally, we discovered that metformin and a potent AMPK activator, PF-06409577, can reverse mitochondrial phenotypic changes in brain endothelial cells and reduce the elevation of VCAM-1 expression associated with chronic CCM disease. Therefore, metformin can provide cytoprotection and may reverse the CCM endothelial phenotype by activating AMPK.

**Conclusions:** Predictions using AI technology and high-throughput drug screening, combined with pharmacokinetic, pharmacodynamic, and toxicological studies in CCM animal models, identified metformin as a promising drug candidate for repurposing for the long-term treatment of CCM disease. We propose that metformin enhances metabolic adaptation to brain vascular malformations by activating AMPK, which helps reverse mitochondrial fragmentation in brain endothelial cells.

**Graphical abstract:** 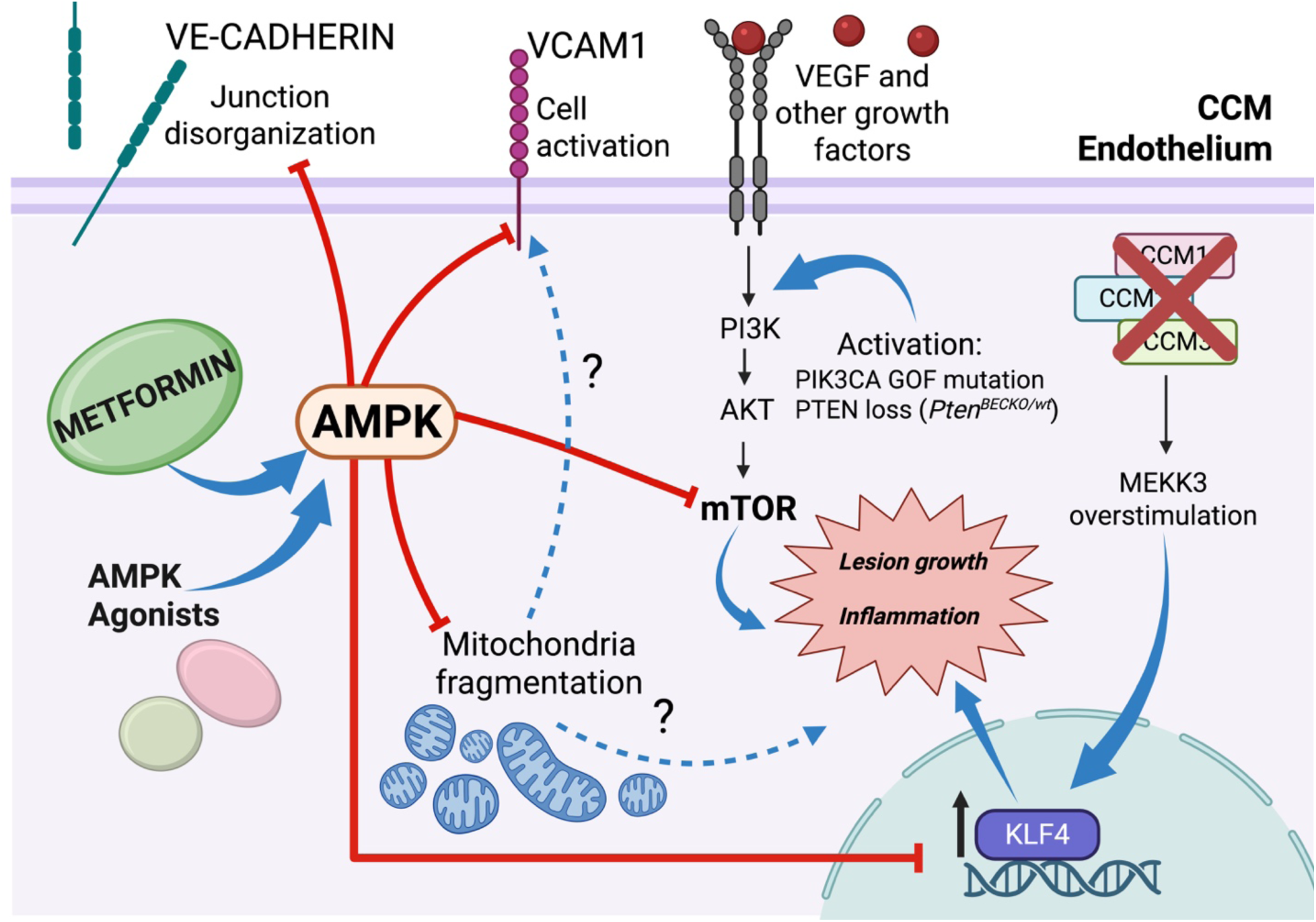

## Introduction

Cerebral cavernous malformations (CCM) are a neurovascular disease that affects the brain and spinal cord, arising mainly due to loss-of-function mutations (which can be inherited germline or somatic mutations) in the genes *KRIT1* (Krev1 interaction trapped gene 1, *CCM1*), *CCM2* (*Malcavernin*), or *PDCD10* (Programmed cell death protein 10, *CCM3*)^1–4^. CCM manifests with various neurological symptoms and lacks effective therapeutic options, only some symptoms are partially alleviated by medication^5–10^. An estimated 0.2% of the US population has a CCM lesion, but only 25% are diagnosed^11,12^. Of these, ∼20% are diagnosed as children (0.0125% of the US population or approximately 40,000 patients)^13^. Around 20% of CCM cases are familial, and approximately 80% are sporadic. Familial CCM is an autosomal dominant disease with variable penetrance, even within the same family^14^. Familial cases usually have multiple lesions, whereas sporadic cases typically have only one lesion. However, at the histological level, the lesions in both sporadic and familial forms of CCM are almost identical^15^. The symptoms most frequently found in patients with symptomatic CCMs are related to motor disability, weakness, seizures, stress, and anxiety^16,17^. The severity of these symptoms may depend on the location of the lesion within the CNS, the stage of the lesions, or the presence of multiple lesions. Life expectancy can be significantly affected by various factors, including mortality from uncontrolled seizures, hemorrhage, or complications from surgery in some instances^16,17^. Patients with the familial form of the condition may require lifelong pharmacological therapy, which is currently not available.

The vascular hypothesis suggests that CCMs develop due to a loss of function of CCM genes within brain endothelial cells^18^. This loss of function results in increased expression of the transcription factors KLF2 and KLF4, which are associated with the onset of CCM pathogenesis^4,19–21^. Alterations in the brain’s vascular structures and functions are known as CCM endothelium^18,22,23^. Recent studies indicate that the CCM endothelium can display various changes depending on the disease^23^ state. These include marked disruption of intercellular junctions^21,24–26^, increased angiogenesis^21,22,27–30^, activation of anti-coagulation mechanisms^4^, neuroinflammation^22,23,31–34^, hypoxia signaling^22,23,35^, and endothelial-mesenchymal transition^36^. There is often an elevation in reactive oxygen species (ROS) levels^1,15,22,37,38^ and increased vascular permeability^24,39–50^. These processes can lead to complications such as hemorrhages^51,52^ and thrombosis^22,23,31,33^, which are significant contributors to morbidity and characteristics of CCM lesions observed in both human cases and CCM animal models^4,21,23,24,35,53,54^. Moreover, the severity of CCM varies due to environmental hypoxia^22^ and genetic factors, including somatic mutations in the PIK3CA (Phosphatidylinositol-4,5-bisphosphate 3-kinase catalytic subunit alpha) and MAP3K3 (Mitogen-Activated Protein Kinase Kinase Kinase 3) genes^7–9,55,56^. Despite several proposed therapies aimed at inhibiting specific signaling pathways affected by CCM disease in preclinical studies, there is currently no effective therapeutic options available for preventing or treating CCM, with surgery being the only viable option^5,54,57^. However, surgical intervention is often unfeasible for lesions in critical or hard-to-reach areas, or for patients with multiple lesions.

Recent advances in AI have revolutionized therapeutic target identification, offering promising applications for complex neurovascular disorders such as CCMs. Using machine-learning models for gene target prioritization in combination with omics-based analysis, the Benevolent AI platform was deployed to identify therapeutic targets for CCM pathology. With supporting evidence from the Benevolent Knowledge Graph^58,59^, potential therapeutic targets were identified, including AMPK and mTOR signaling, to improve the altered metabolic and transcriptomic processes observed in CCM endothelium. A high-content screening of approximately 100 FDA-approved drugs identified metformin as an AMPK agonist and mTOR inhibitor. Metformin was found to reverse alterations in cell-cell junction organization and increase transcription factor KLF4 in human CCM endothelial cells in cultured assays. Preclinical studies showed that metformin can effectively prevent the progression of CCM disease by preventing the development of multiple cavernous lesions in the brains of CCM mouse models. The protective effect was linked to increased AMPK activity due to enhanced phosphorylation of AMPK and reduced mTOR signaling, as shown by decreased phosphorylation of S6 ribosomal protein, a pharmacodynamic markers. Furthermore, our toxicological studies on CCM animal models indicate that long-term use of therapeutic dose of metformin is associated with a safe profile. We demonstrate that metformin and the AMPK agonist PF-06409577 reverse mitochondrial phenotypic changes and the inflammatory marker VCAM-1 in brain endothelial cells derived from chronic CCM mouse models. In this study, AI-based technology has identified a combination of elevated AMPK levels and modulation of mTOR signaling as therapeutic approaches for CCM pathology. We have established the preclinical safety and efficacy of a pharmacological approach that could expedite clinical trials by repurposing metformin for the long-term treatment of CCM disease. This approach may influence metabolic adaptation to brain vascular malformations by activating AMPK and regulating mitochondrial dynamics.

## Material and methods

### Artificial intelligence target predictions

Targets were predicted from the Benevolent Knowledge Graph (the Benevolent Platform™)^60^. At the core of the platform lies a comprehensive, disease-agnostic Knowledge Graph that synthesizes diverse biomedical information from multiple independent sources. This includes data extracted from more than 35 million scientific publications and numerous structured databases. The platform employs machine learning models to identify and analyze complex relationships between biomedical concepts within the Knowledge Graph. The target predictions were prioritized based on their plausibility to moderate CCM biology, novelty for the indication, safety, and expression in relevant cell types, using data surfaced from the Knowledge graph.

### Cell culture

Human brain endothelial cell line, hCMEC/D3, at passages 23–33, was routinely cultured in EGM-2 MV medium (Lonza), hereafter referred to as EGM-2 medium. This medium was supplemented with the following components obtained from the manufacturer at specified concentrations: 0.025% (v/v) rhEGF, 0.025% (v/v) VEGF, 0.025% (v/v) IGF, 0.1% (v/v) rhFGF, 0.1% (v/v) gentamicin, 0.1% (v/v) ascorbic acid, 0.04% (v/v) hydrocortisone, and 2.5% (v/v) FBS^61,62^. Tissue culture flasks were precoated with a 1:20 solution of collagen type I. The cells were then seeded onto the collagen-coated flasks and maintained at 37°C in a 95% air and 5% CO2 atmosphere until confluence.

### Human CCM in vitro model

We adopted an inducible RNAi system (to knockdown CCM genes) that allows us to identify retrovirally transduced cells and RNAi induction through the expression of two fluorescent reporters, as previously reported^63^. This approach uses an inducible tetracycline-responsive element (TRE) promoter that controls the expression of a dsRed fluorescent protein and a microRNA-embedded shRNA targeting *PDCD10* and a second promoter, the phosphoglycerate kinase (PGK), that controls the constitutive expression of the yellow-green fluorescent protein Venus^63^ which we denominated TRMPV-*PDCD10* (TRE-dsRed-miR30-against-PDCD10-PGK-Venus). We generated stable hCMEC/D3 cell lines using this RNAi system, each expressing one of three different TRMPV-*PDCD10* constructs. We observed that hCMEC/D3 cells transduced with TRMPV-PDCD10 retroviral particles, purified by FACS, and cultured in the presence of 10 μg/mL doxycycline for 20 days produced cells with 98% TRE-dsRed-miR30-against-PDCD10-PGK-Venus-positive.

### Drug screening in human CCM in vitro model

For drug screening experiments, 2.5 x10^5^ cells/cm^2^ were plated per well on collagen-precoated Milicell EZ slide 8-well glass (PEZGS0816, Millipore Sigma) for 48 hours. All FDA-approved drugs were prepared at a concentration of 10 mM in DMSO. Then, drugs were diluted in a reduced EGM-2 medium (0.012% (v/v) rhEGF, 0.012% (v/v) IGF, 0.05% (v/v) rhFGF, 0.1% (v/v) gentamicin, 0.1% (v/v) ascorbic acid, and 2.5% (v/v) FBS) and tested at three different concentrations (10 µM, 1 µM, and 0.1 µM) for 72 hours with a media change at 48 hours. Cells were then processed for immunohistochemistry.

### Genetically modified animals

Brain endothelial cell (BEC)-specific conditional *Pdcd10*-knockout mice were generated by crossing a brain endothelial tamoxifen-regulated Cre recombinase (*Slco1c1-iCreERT2*) strain with loxP-flanked *Pdcd10* (*Slco1c1-iCreERT2; Pdcd10^fl/fl^*). On postnatal day 3 (P3), mice were administered 50 μg of 4-hydroxy-tamoxifen by intragastric injection to induce inactivation of the endothelial *Pdcd10* gene. Conditional *Pdcd10*-knockout mice were also crossed with a MITO-tag strain (*B6.Cg-Gt(ROSA)26Sor^tm1(CAG-EGFP)Brsy^/J*) to obtain a mouse model that conditionally expresses, in a Cre recombinase-dependent manner, a hemagglutinin (HA)-tagged EGFP that localized to mitochondrial outer membrane (*Slco1c1-iCreERT2; Pdcd10^fl/fl^; B6.Cg-Gt(ROSA)26Sor^tm1(CAG-EGFP)Brsy^/J, [Pdcd10^BECKO^;MITO-tag]).* On postnatal day 1 (P1), mice were administered 50 μg of 4-hydroxy-tamoxifen by intragastric injection to induce inactivation of the *Pdcd10* gene and the expression of the HA-tagged EGFP in brain endothelial cells. As controls, littermates with no floxed copies of *Pdcd10* were used (*Slco1c1-iCreERT2; Pdcd10^wt/wt^; B6.Cg-Gt(ROSA)26Sor^tm1(CAG-EGFP)Brsy^/J, [Pdcd10^wt/wt^;MITO-tag]*). Additionally, BEC-specific conditional *Krit1*-knockout mice were generated by crossing our *Slco1c1-CreERT2* strain with a LoxP-flanked *Krit1* (*Slco1c1-CreERT2; Krit1^fl/fl^*) strain. The *Slco1c1-CreERT2; Krit1^fl/fl^* strain was then crossed with a *Pten-knockout* strain (*Slco1c1-CreERT2; Krit1^fl/fl^*; *Pten^tm1Hwu/J^*) to generate a model of enhanced PI3K-Akt-mTOR activity in CCMs. On postnatal day 7 (P7), mice were administered 30 μg of tamoxifen by intragastric injection to induce inactivation of the *Krit1* and *Pten* genes in brain endothelial cells. All animals were kept in a pathogen-free facility and were housed in groups of 5 per cage under a standard light cycle (12 h:12 h light:dark). The main mouse sub-strain used in these studies was *C57BL/6J,* and all mice were provided access to food and water *ad libitum*. All animal experiments were performed at the University of California, San Diego, and in compliance with animal procedure protocols approved by the University of California, San Diego Institutional Animal Care and Use Committee.

### Metformin intraperitoneal injections

For *Slco1c1-CreERT2; Krit1^fl/fl^*; *Pten^tm1Hwu/J^* mice were injected intraperitoneally (IP) with metformin (50 ug/g) once daily beginning on postnatal day 14 (P14) with a total volume of 100 ul and 50 ul for the next consecutive days until their last injection at P18 where brains were collected for histological analysis. For strains *Slco1c1-CreERT2;Pdcd10^fl/fl^,* mice were injected IP with metformin (100 mg/kg) with a total volume of 100 ul once every three days at P30 until their last injection at P60 where brains were collected for histological analysis. Control groups were injected IP with saline on its own.

### Immunofluorescence

For the drug screening, cells grown in precoated Milicell EZ slide 8-well glass (PEZGS0816, Millipore Sigma) were fixed in 4% paraformaldehyde (PFA) at room temperature (RT) for 10 min, followed by permeabilization with 0.5% Triton X-100 in PBS for 10 min at RT, and blocked with BSA 0.5% in PBS for 1hr. Cells were then incubated in rabbit monoclonal antibodies against VE-cadherin (1:200; 2500S; Cell Signaling) and goat polyclonal antibodies against KLF4 (1:100, AF3640; R&D Systems) at 4°C overnight. For mitochondrial dynamics analysis, cells were incubated with rabbit monoclonal antibodies against HA-tag (1:500, 3724S; Cell Signaling), and rat monoclonal antibodies against VCAM-1/CD106 (1:100; 105701; BioLegend). After incubation with primary antibodies, cells were washed three times in PBS and incubated in suitable Alexa Fluor coupled secondary antibodies (1:300, Jackson Laboratory) in PBS for 2h at RT. Cell nuclei were stained with DAPI, and slides were mounted using Fluoromount-G mounting medium (SouthernBiotech).

Brains from *Pdcd10^BECKO^* mice at the specific age were isolated and fixed in 4% PFA in PBS at 4°C overnight. Tissue was cryoprotected in 30% sucrose solution in PBS, and then embedded and frozen in O.C.T compound (Fischer Scientific). Brains were then cut using a cryostat into 16-µm sagittal sections using a cryostat and mounted onto Superfrost Plus slides (VWR International). Sections were incubated in a blocking-permeabilization solution (0.5% Triton X-100, 5% goat serum, 0.5% BSA, in PBS) for 2 hours and incubated in rat polyclonal antibodies against CD31/PECAM1 (1:100; AF3628; R&D Systems), rabbit polyclonal antibodies against pS6 (1:200; 4858S; Cell Signaling), rabbit polyclonal antibodies against pAMPK (1:250; 2531S; Cell Signaling), rabbit monoclonal antibodies against HA-tag (1:100, 3724S; Cell Signaling) in PBS at room temperature overnight. When staining for isolectin-B4 (IB4), slides were incubated with IB4 biotin-conjugated (1:80; L2140; Sigma-Aldrich) in Pblec buffer (1X PBS, 1mM CaCl_2_, 1mM MgCl2, 0.1 mM MnCl_2_, and 0.1% Triton X-100) overnight followed by incubation with streptavidin (Alexa-Fluor 647; 1:100; S21374; Invitrogen) in Pblec buffer for 24 hrs at RT. Tissue sections were washed three times in PBS and incubated with suitable Alexa Fluor-coupled secondary antibodies (1:300, Jackson Laboratory) in PBS for 2h at RT. Cell nuclei were stained with DAPI, and slides were mounted using Fluoromount-G mounting medium (SouthernBiotech).

For cells in the drug screening and mouse tissue samples, the slides were viewed with a high-resolution slide scanner (Olympus VS200 Slide Scanner), and the images were captured with VS200 ASW V3.3 software (Olympus). For mitochondria analysis, slides were viewed with a SP8 Confocal with Lightning Deconvolution (Leica). Quantifications were performed blinded. The quantification analysis was performed using ImageJ Ver. 1.53f on high-resolution images. Four to five brain sections per mouse were used for analysis. Twenty to twenty-five cells were quantified for the mitochondrial analysis per condition.

### VE-cadherin junction analysis

VE-cadherin junction was quantified using the Junction Analyzer Program ^64^ following the developer User-Guide available at https://github.com/StrokaLab/JAnaP. Briefly, cells whose perimeters were visible in each image were traced via “waypointing” in the software. Waypointing was performed using VE-cadherin staining, using 150 cells per treatment. The different cell junction characteristics (coverage, continuous, discontinuous, and punctate) were quantified for each cell as a percentage of the cell edge.

### KLF4 nuclear quantification

Nuclear KLF4 staining was done using ImageJ2 (Fiji) ^65^ as previously reported^66^. Nuclei images stained with DAPI were transformed into binary images to define the regions to be quantified for KLF4 using the AnalyzeParticles function.

### Mitochondrial morphology and network analysis

Microvascular brain endothelial cells were isolated from *Pdcd10^BECKO^; MITO-tag* and control *Pdcd10^wt/wt^; MITO-tag* mice (P70) and maintained in cultured conditions for 5 days. On the sixth day after isolation, cells were treated with Metformin (50 µM; Selleckchem, S1950) or vehicle, and AMPK agonist PF-06409577 (0.1 M; Selleckchem; S8335) or vehicle, and maintained for 72 hrs for either treatment. After treatment, cells were fixed with 4% PFA and analyzed for mitochondrial dynamics by immunofluorescence. The ImageJ pipeline used followed the protocol published by Chaudhry et al^67^. Mitochondrial features analyzed include area, perimeter, shape parameters (i.e., aspect ratio, form factor) and network complexity (i.e., branches, branch junctions, branch length).

### RNA isolation

Total RNA from brain endothelial cells was isolated by TRIzol method according to the manufacturer’s instructions (Thermo Fisher Scientific). The lysates were transferred to Phase Lock Gel 2ml tubes, and 200 µL of chloroform (Thermo Fisher Scientific) was added to each tube, mixed vigorously for 15 seconds, and incubated at room temperature for 3 minutes. Samples were centrifuged at 12000g for 10 minutes at 4°C, and aqueous phases were collected and transferred into 1.5ml DNAse/RNAse free tubes. For RNA precipitation, 500µl of isopropanol was added, and samples were resuspended and incubated for 10 minutes at room temperature, followed by centrifugation at 12000g for 10 minutes at 4°C. After supernatant removal, the pellet was washed with 1ml of 75% ethanol, followed by centrifugation at 7500g for 5 minutes at 4C. Afterward, RNA was resuspended in RNAse-free water, and the quantity (ND-1000 spectrophotometer; NanoDrop Technologies) and quality (TapeStation; Agilent) of total RNA were analyzed.

### Histological analysis and lesion quantification

Brains from *Krit1^BECKO^;Pten^BECKO/wt^* mice at P18 and *Pdcd10^BECKO^* animals at P60, injected with either Metformin or vehicle, were fixed in 4% PFA overnight. After cryoprotection in sucrose (30%) and freezing, 16 μm sections of sagittal brain tissue were cut onto Superfrost Plus slides (VWR International), and stained following the same orcein and martius scarlet blue (OMSB)^22^ protocol. For OMSB, brain sections were post fixed in Bouin’s solution at 56°C for 1 h, then rinsed for 10 min in tap water to remove traces of picric acid. Staining with orcein was performed for 20 min at 56°C, then slides were incubated in acidic alcohol for 2 min, and transferred to 96% ethanol for 30 s. Sections were then stained with martius yellow for 3 min and rinsed in distilled water (2×2 min), followed by staining with crystal scarlet for 10 min. Differentiation with phosphotungstic acid for 2 min was followed by a 10 sec in distilled water. Sections were then stained with methyl blue for 5 min and rinsed in acetic acid water for 3 min. Dehydration through ethanol solutions of increasing grade was performed, and then sections were cleared in xylene and mounted in Cytoseal XYL (Epredia). Stained sections were imaged using NanoZoomer Slide Scanner (Hamamatsu Photonics; San Diego, USA). Quantification of CCM lesions was performed in a blinded manner. The quantification analysis was performed using Hamamatsu Photonics software. Lesion area and lesion number per brain section area were calculated.

### Statistical analysis

Statistical analyses for in vitro data and visualizations were performed using R (version 4.1). For each morphological or molecular parameter VE-cadherin coverage, continuous, punctate, and discontinuous, and KLF4 nuclear intensity descriptive statistics, including mean, standard deviation, and median, were calculated within each treatment group. To quantify how each treatment restored endothelial junction morphology or KLF4 nuclear localization to levels comparable to PDCD10-WT (control) levels, we developed a *recovery score* metric. This score was determined independently for VE-cadherin staining (using four parameters: coverage, continuity, discontinuity and punctation) and for nuclear KLF4 staining (using integrated density). Raw single-cell measurements for all parameters were merged into a single long-format table linking each cell to its corresponding treatment and concentration.

For every parameter, the median value was computed for the siPDCD10 (TRMPV with doxycycline) and control (TRMPV-PDCD10 without doxycycline) groups and used as reference anchors. Each parameter was also assigned a direction of improvement: parameters with higher values indicating better morphology (coverage, continuous) were classified as “high”, whereas those with lower values indicating improved morphology (punctate and discontinuous) were classified as “low”.

For each cell (i) and parameter (p), an individual recovery score was computed as:

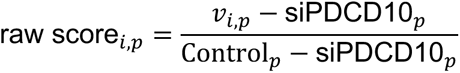

Where 𝑣_𝑖,𝑝_is the observed value for the given drug, concentration, and parameter, and Control_𝑝_ and siPDCD10_𝑝_are the respective medians for the control and siPDCD10 reference groups. To maintain consistent directionality across parameters, values for parameters where “low” indicates improvement were sign-inverted prior to calculation. The resulting *raw scores* were then clipped to the range [0-1], such that: 0 represents a PDCD10-KD-like phenotype, 1 represents a PDCD10-WT-like phenotype, and intermediate values represent partial recovery. The same approach was applied to KLF4 nuclear staining, with the Integrated nuclear density (IntDen) used as a quantitative measure, in this case, lower nuclear KLF4 was a sign of recovery. To ensure robust interpretation, raw data were assessed for normality using the Shapiro-Wilk test for each parameter and treatment group, with results categorized as “normal” or “not normal” based on a p-value threshold of 0.05. Because the data did not meet normality assumptions, the nonparametric Kruskal-Wallis test was used to compare each parameter across all treatment groups. Pairwise comparisons were subsequently conducted using Dunn’s test with Bonferroni correction to identify specific group differences, with adjusted p-values reported (*p* < 0.05, *p* < 0.01, *p* < 0.001).

For visualization, heatmaps and boxplots were generated using ggplot2 to depict the distribution of each parameter across treatment groups. In heatmaps, lighter colors indicate increased recovery, while darker colors indicate no recovery. For boxplots, horizontal dashed lines representing median values for control reference conditions (siPDCD10 and control) were included in each plot, colored red and blue, respectively, to highlight baseline differences.

Statistical analysis for in vivo experiments was performed using Prism software (GraphPad). Data are expressed as mean values ± standard error of the mean (SEM) for multiple individual and biological experiments. For all experiments, the number of independent and biological replicates (n) is indicated. Mice experiments were randomized, and histological quantification was performed blinded. The Shapiro-Wilk normality test with alpha=0.05 assessed the normality of the data. For comparison between two groups, Student’s unpaired two-tailed *t-test* or analysis of variance (ANOVA) for comparison among multiple groups, followed by the Tukey post hoc test, was used for normally distributed data. The Mann-Whitney two-tailed test was used for two groups, and the Kruskal-Wallis test, followed by Dunn’s post hoc test, was used for non-normally distributed data. Sample sizes were calculated by assuming a two-sided alpha=0.05, 80% power, and homogeneous variances for two samples with the means and standard deviation from previous studies^4,13,15,16^. At least three biologically independent experiments were conducted to ensure reproducibility. No statistical across-test comparisons were performed in this manuscript.

## Results

### Artificial intelligence (AI) and high-content screening identify pharmacological targets for CCM disease

BenevolentAI’s machine learning algorithms combine diverse data types from various biomedical entities, including diseases, genes, tissues, and biological mechanisms. This integration, along with our previous transcriptomics data derived from CCM models^15,21,35^, enable the prioritization of potential therapeutic drug targets for CCM disease. We then triaged and selected targets that meet acceptable safety profiles (Fig. 1A; Supplemental Table 1). A list of targets was selected for experimental validation, using a human CCM cell culture model by generating a tetracycline-regulated RNAi directed^63^ against the CCM3 (*PDCD10*) mRNA transcript in a human brain endothelial cell line (hCMEC/D3). The construct uses an inducible tetracycline-responsive element (TRE) promoter that controls the expression of a dsRed fluorescent protein and a microRNA-embedded shRNA directed against *PDCD10* (CCM3 gene) and a second promoter, the phosphoglycerate kinase (PGK), that controls the constitutive expression of the yellow-green fluorescent protein Venus^63^ that we designated TRMPV-*PDCD10* (TRE-dsRed-miR30-against-PDCD10-PGK-Venus)^18^ (Fig. 1B). After fifteen days, doxycycline-treated cells (SiPDCD10) exhibited significant disruption of VE-cadherin staining in the junctions and altered gene expression characteristics of CCM disease^15,18,21,23,35,68,69^ compared to vehicle-treated cells. This model was then used to perform high-content screening analysis to assess drug molecules that could reverse alterations in cell-cell junction organization and the elevation of KLF4 in CCM endothelial cells, which are strong markers of CCM phenotypes in cultured cells^21^. We conducted immunofluorescence analysis using antibodies specific to VE-cadherin and the transcription factor KLF4 (Fig. 1C,1D,1E). We tested pharmacological drugs at concentrations of 0.1 μM, 1 μM, and 10 μM to determine the effective dose. Brain endothelial cells treated with doxycycline, (SiPDCD10 cells) were exposed to inhibitors or agonists for 72 hours, along with control cells (TRMPV-PDCD10), that were not treated with doxycycline. High-content screening validation identified drugs targeting CAMKK1/2, KEAP1, ERBB2, KDR, KAT2B/2A, and IKBKB as inhibitors that significantly reversed VE-cadherin staining and normalized KLF4 expression levels in siPDC10 brain endothelial cells (human CCM model) at most tested doses. We found that several pharmacological drugs significantly normalized KLF4 expression levels without significantly affecting the reversal of VE-cadherin staining in siPDC10 brain endothelial cells. These drugs included inhibitors targeting KRAS, RAF1/BRAF, PPARG, PRKDC, IKBKB, MBTPS1, CAMK2B/2A, NLRP3, INPPL1, ADAM10, PLAU, LYN, CTSC, MIF, and NTRK2. Notably, we also identified pharmacological inhibitors with rescue effects at low doses, including CFTR, BRAF, FLT3/PDGFRA/B, and mTOR, among others. High-content screening and dosing optimization identified metformin, a known AMPK agonist and an mTOR inhibitor, as an effective drug capable of reversing changes in cell-cell junction organization and reducing the increase of the transcription factor KLF4 in human CCM endothelial cells during culture assays (Fig. 1D,1E). Notably, the AMPK agonist (PF-06409577), but not the mTOR inhibitor (AZD-8055), was effective in reversing these alterations in cell-cell junction organization in human CCM endothelial cells at the doses tested (Supplemental Table 1). We further chose metformin for preclinical CCM studies based on metformin’s favorable safety profiles in children and adults^70,71^, a wide therapeutic window, tolerable doses (range from 500 mg/day to 2000 mg/day)^72,73^, direct effects on endothelium^74–76^, and well-characterized pharmacokinetics and tissue distribution studies (including in brain tissue)^77,78^.

**Figure 1.**
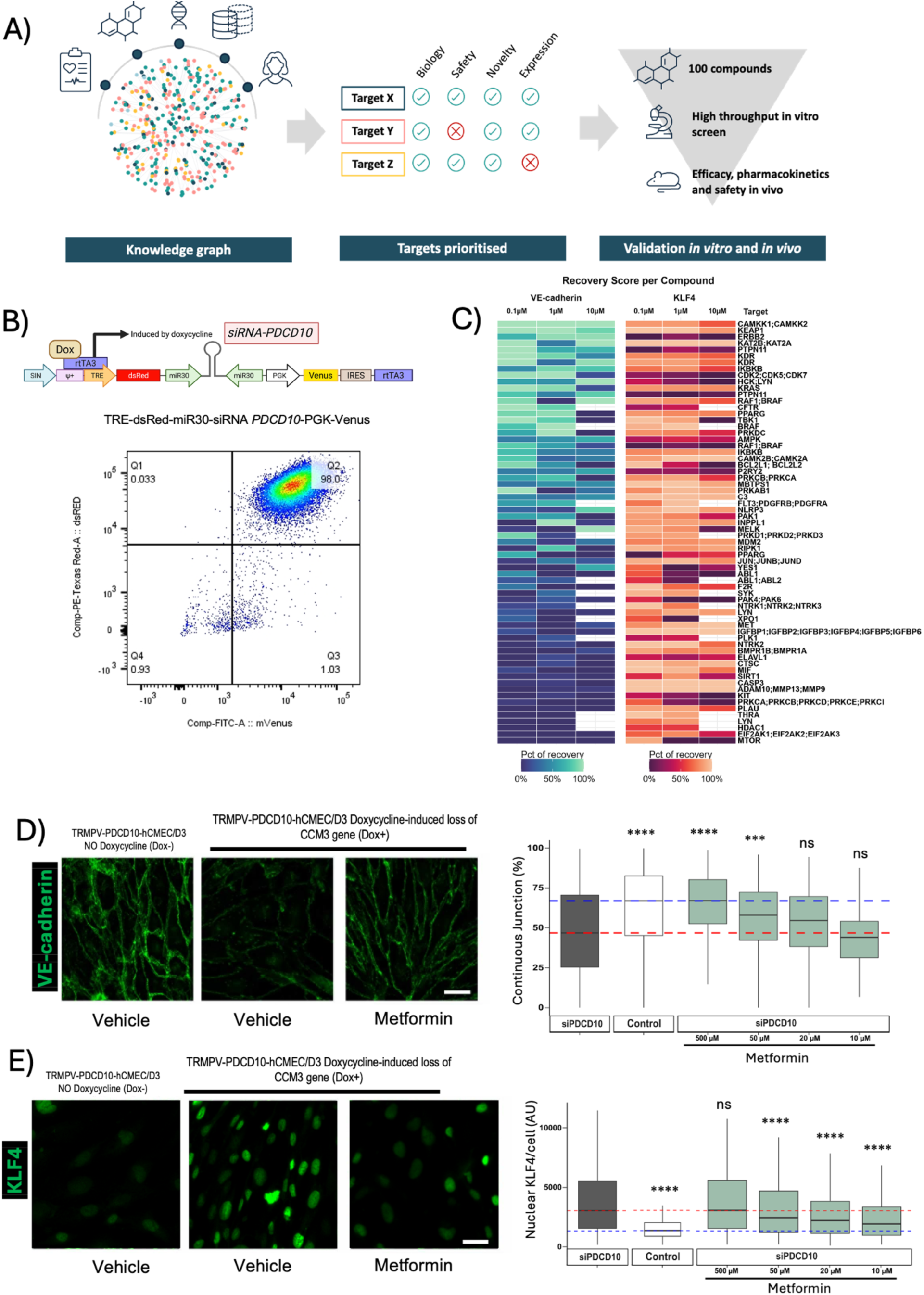
AI and high-content screening assay identify pharmacological targets for CCM disease. **A)** Schematic of Benevolent Platform^TM^ AI methodology applied to target identification/validation, with a knowledge graph composed of many data modalities, including biomedical literature, transcriptomic, and clinical data to identify therapeutic targets for CCMs. These targets were assessed and prioritised based on biological rationale, novelty, and safety, prior to testing in a high-throughput in vitro screen. After selecting targets with suitable FAD-approved drugs, we progressed to in vivo models for efficacy and pharmacokinetic testing. **B)** Top: schematic of the RNAi system for PDCD10 (TRMPV-PDCD10) showing the inducible tetracycline-responsive element (TRE) that regulates the expression of the dsRed fluorescent protein and a microRNA-embedded shRNA targeting PDCD10, followed by a phosphoglycerate kinase (PGK) promoter that drives constitutive expression of the Venus fluorescent protein. Bottom: Representative flow cytometry plots of transduced hCMEC cells with TRMPV-PDCD10 treated with doxycycline. The plot shows the percentage of Venus-positive (indicating transduction) and dsRed-fluorescent (confirming expression of the microRNA-embedded shRNA). **C)** Heatmaps displaying the percentage of recovery for VE-cadherin staining (left) and KLF4 nuclear staining (right) in siPDCD10 treated with various compounds at concentrations of 0.1, 1, and 10 μM. Color scales indicate the recovery percentage. Concentration is shown in each column. The right annotation indicates target genes. **D)** Left: VE-cadherin immunostaining (green) in control cells (no doxycycline), and siPDCD10 cells (+ Doxycycline) treated or not with metformin. Right: quantification of VE-cadherin continuous junction shown as percentage (%) for siPDCD10 and control cells, along with siPDCD10 treated with decreasing concentrations (500, 50, 20, and 10 μM) of metformin. Pairwise comparisons were conducted using Dunn’s test with Bonferroni correction; significance corresponds to adjusted p-values (ns, no significant; *, *p* < 0.05; **, *p* < 0.01; ***, *p* < 0.001; ****, *p* < 0.001). **E)** Left: KLF4 immunostaining (green) in control cells (no doxycycline), and siPDCD10 cells (+ Doxycycline) treated or not with metformin. Right: quantification of nuclear KLF4/cell shown as arbitrary units (AU) for siPDCD10 and control cells, along with siPDCD10 treated with decreasing concentrations (500, 50, 20, and 10 μM) of metformin. Pairwise comparisons were conducted using Dunn’s test with Bonferroni correction; significance corresponds to adjusted p-values (ns, no significant; *, *p* < 0.05; **, *p* < 0.01; ***, *p* < 0.001; ****, *p* < 0.001).

### Metformin treatment reduces neurovascular lesions in an aggressive CCM1 mouse model

The varying severity levels observed among patients with CCM^5–10^ may be linked to the presence of somatic PIK3CA (Phosphatidylinositol-4,5-Bisphosphate 3-Kinase Catalytic Subunit Alpha) activating mutations^7–9,55^. Recent studies have shown that CCM lesions with PIK3CA activating mutations account for 60-70% of surgically removed CCMs. This suggests that the PIK3CA gain-of-function mutation contributes to increased growth and bleeding in CCM lesions, often necessitating surgical intervention^7–9,55^. To investigate whether metformin could be used as therapeutic option for aggressive CCM1 lesions in need of surgery, we utilized an aggressive animal model of CCM1 by crossing a model of brain endothelial deletion of *Krit1* (which represents CCM1, the most common form of the disease) with deletion of one copy of of the *Pten* gene (phosphatase and tensin homolog). This combination simulates CCM1^23^ disease and a “gain of function”^9,79^ in PI3K by enhancing PI3K-mediated activation of the canonical AKT-mTOR pathway due to the loss of PTEN activity. The resulting model develops rapidly growing CCM lesions (*Slco1c1-CreERT2;Pdcd10^fl/fl^;Pten^fl/wt^* or *Krit1^BECKO^;Pten^BECKO/wt^*) (Fig. 2). We observed that daily intraperitoneal injections (IP) of metformin at a dose of 50 mg/kg/day were well tolerated in both male and female *Krit1^BECKO^;Pten^BECKO/wt^* mice from postnatal day (P) 12 to 20. However, increasing the dosage to 100 mg/kg/day resulted in lethality in this neonatal model of CCM disease (data not shown). Therefore, we conducted experiments in which *Krit1^BECKO^;Pten^BECKO/wt^* animals were injected intraperitoneally with either metformin (50 mg/kg/day) or a saline solution as control vehicle for five consecutive days, from P14 to P18 (Fig. 2A,2B). Since metformin is known to activate AMPK and inhibit mTOR activity, we first conducted immunostaining for phosphorylated AMPK (pAMPK) to assess its activation. To evaluate mTOR activity inhibition, we performed immunostaining for the phosphorylated S6 ribosomal protein (pS6), as its phosphorylation is dependent on mTORC1 activity, both serving as a pharmacodynamic markers of metformin. Histological analysis showed increased pAMPK and decreased pS6 immunofluorescent signal in CCM lesions of *Krit1^BECKO^;Pten^BECKO/wt^* animals treated with metformin when compared to CCM lesions of *Krit1^BECKO^;Pten^BECKO/wt^* animals treated with vehicle (Fig. 2C). Using orcein staining, we conducted a blinded quantitative analysis of CCM lesions. This analysis revealed a significant decrease in the area of multi-cavernous lesions in *Krit1^BECKO^;Pten^BECKO/wt^* mice treated with metformin, compared to those injected with a saline vehicle (Fig. 2C-2H). However, the histological analysis did not show any significant changes in single CCM lesions. These results suggest that the FDA-approved drug metformin, which acts as an AMPK agonist and mTOR inhibitor, may represent a potential treatment for rapidly growing multicavernous CCM lesions.

**Figure 2.**
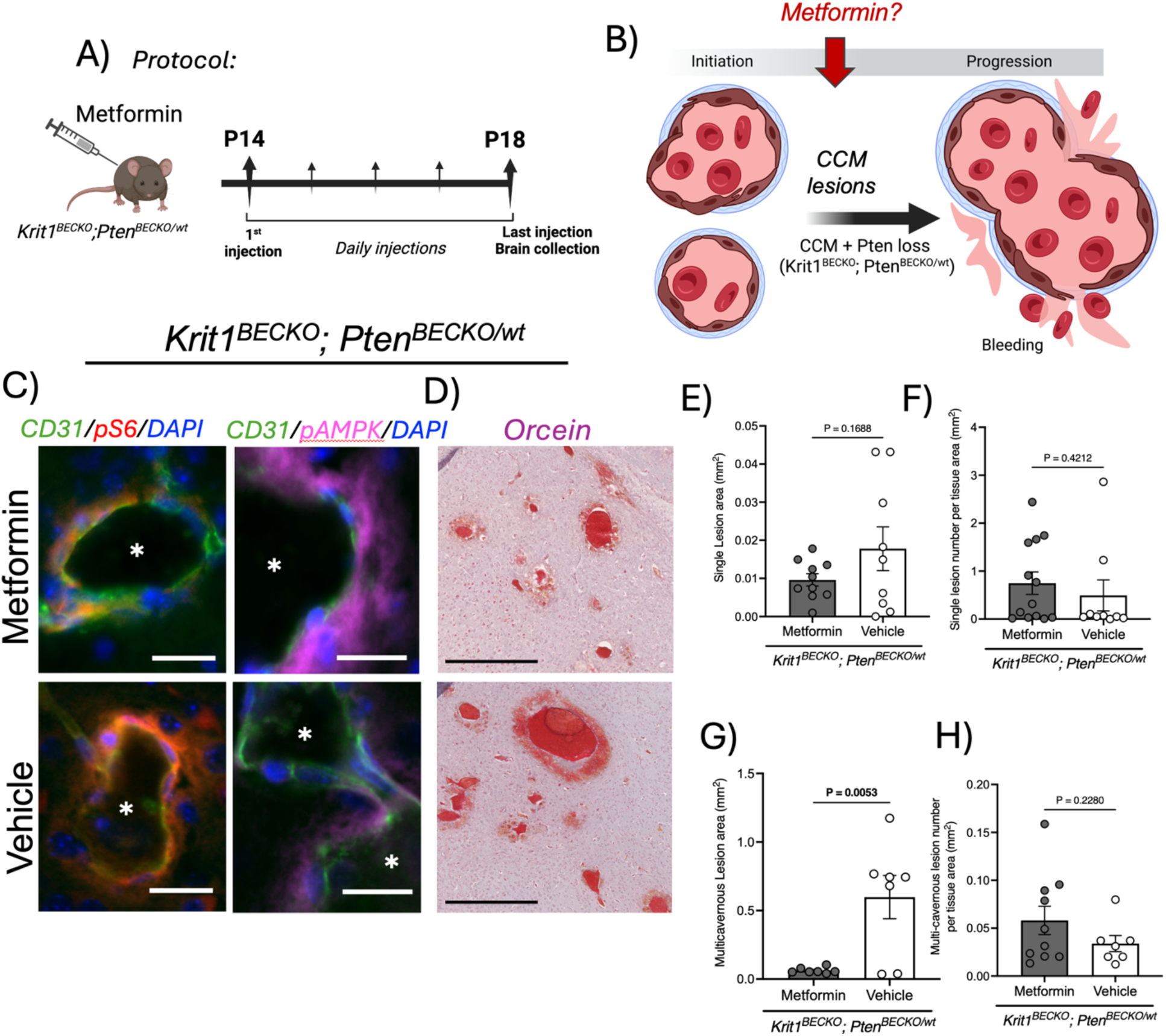
Metformin reduces lesion burden in an acute mouse model of CCM. **A)** Protocol of intraperitoneal injections of Metformin (50 µg/g) or vehicle in *Krit1^BECKO^; Pten^BECKO/wt^* mice (P14). Metformin was injected daily for 5 days and brains were collected at P18. **B)** Hypothesis on the effect of metformin on acute CCMs. Histological analysis of *Krit1^BECKO^; Pten^BECKO/wt^* mouse brains treated with Metformin versus vehicle. **C,** Immunofluorescence analysis shows decreased pS6 and increased pAMPK expression in animals treated with Metformin in comparison to animals treated with vehicle. **D)** Orcein staining shows presence of bleeding CCM lesions throughout the CNS in both treatment groups. **E-F)** Analysis in single CCM lesion area or number. **G-H)** Analysis in multi-cavernous CCM lesions in area and numbers. N= 7 or 10 mice per group.

### Metformin treatment reduces neurovascular lesions in chronic *Pdcd10^BECKO^* mice

We previously reported a mouse model that develops a chronic CCM3 disease that closely mimics the phenotype observed in humans with CCMs^4,23,31,35,80–82^. Therefore, we used this model to assess the preclinical safety and efficacy of long-term metformin therapy for treating CCM disease. This model used *Pdcd10^fl/fl^* mice that were crossed with the brain endothelial tamoxifen-regulated Cre (Slco1c1-CreERT2) to produce *Slco1c1-CreERT2;Pdcd10^fl/fl^*, a “brain endothelial-specific *Pdcd10* knockout” (*Pdcd10^BECKO^*, CCM3)^35,80,82,83^. The chronic murine CCM model involved the gene deletion of the brain endothelial *Pdcd10* in both male and female mice through intragastric injection of 50 ug of 4-hydroxy-tamoxifen in sunflower oil at P6, producing *Pdcd10^BECKO^* mice in which lesions can be monitored at a chronic stage by P60^23,82^. Moreover, while metformin’s pharmacokinetics and brain exposure have been investigated in humans and various species, including mice^77,78^, further studies are necessary to ensure its safe application as a pharmacological treatment for CCM disease (Fig. 3). We, therefore, first observed that metformin, administered at a dose of 100 mg/kg/day, was well tolerated by both male and female P30 *Pdcd10^BECKO^* mice over 20 consecutive days. We did not observe toxicity, as evaluated by complete blood count, metabolic panel analysis (Supplemental Figure 1), and T-cell analysis from the spleen, and histology analysis (data not shown). We also observed that a single IP injection of 100 mg/kg resulted in a plasma metformin concentration (C_max_) of 44.2 µg/ml, reached almost immediately, with a time to maximum concentration (T_max_) of 0.25 hours. After 2 hours, 50% of the maximum metformin concentration remained (t1/2 of 2 hours), and the concentration steadily decreased over the following 8 hours (Fig. 3A). We then conducted experiments to evaluate the efficacy of long-term metformin therapy in *Pdcd10^BECKO^* mice. Specifically, P30 *Pdcd10^BECKO^* mice were injected with either metformin (100 mg/kg) or a saline solution as a vehicle every 3 days for a total of 30 days (Fig. 3B, 3C). We observed that, 30 minutes after the last IP injection of metformin following a long-term therapeutic regimen, the metformin exposure in the *Pdcd10^BECKO^* brains reaches approximately 200 ng/g. Histological analysis of pharmacodynamic markers indicated that CCM lesions in *Pdcd10^BECKO^* brains exhibited increased levels of pAMPK and decreased levels of pS6 in mice treated with metformin compared to those treated with vehicle (Fig. 3E). In addition, quantitative analysis of CCM lesions in the brains of *Pdcd10^BECKO^* mice treated with metformin revealed a significant reduction in the number of multi-cavernous lesions compared to *Pdcd10^BECKO^* mice that received vehicle injections (Fig. 3F-3J). However, we did not observe any significant differences in the lesions or in single CCM lesions between the two groups (Fig. 3G-I). Overall, these studies suggest that metformin could potentially be repurposed as a pharmacological treatment for long-term treatment of CCM disease. The findings also suggest that metformin may be particularly effective in preventing the progression from single lesions to multi-cavernous lesions in CCM by activating endothelial AMPK and lowering mTOR activity.

**Figure 3.**
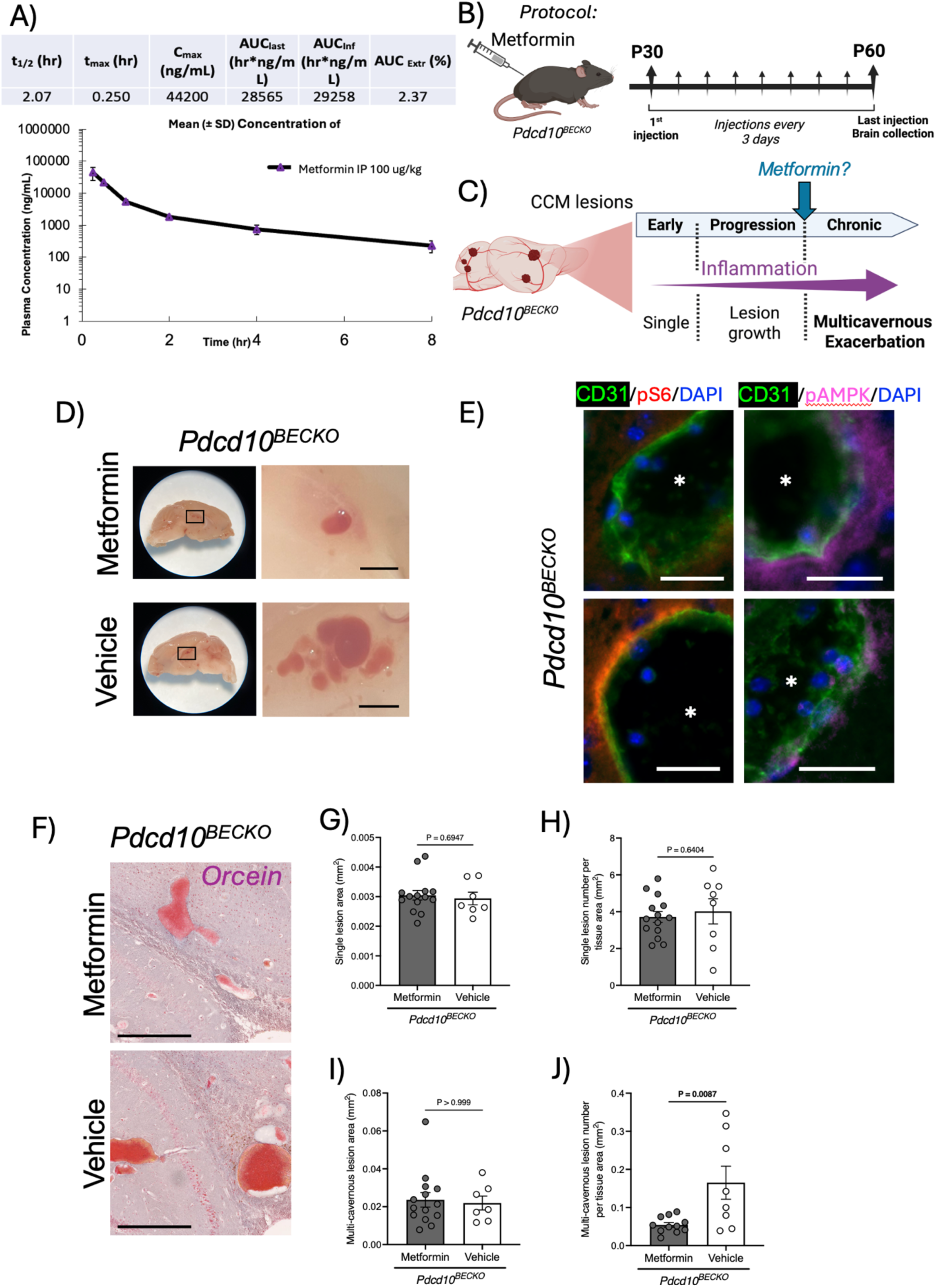
Metformin reduces lesion burden in a chronic mouse model of CCM. **A)** Pharmacokinetic analysis of Metformin in plasma of *Pdcd10^BECKO^* mice (P30). **B)** Protocol of intraperitoneal injections of Metformin (100 µg/g) or vehicle in *Pdcd10^BECKO^* mice (P30). Metformin was injected every 3 days for 30 days and brains were collected at P60. **C)** Hypothesis on the effect of metformin on chronic CCMs. **D)** Macroscopic images of CCM lesions in *Pdcd10^BECKO^* mice injected with Metformin or vehicle. Histological analysis of *Pdcd10^BECKO^* brains from mice treated with Metformin versus vehicle. **E,** Immunofluorescence analysis shows decreased pS6 and increased pAMPK expression in animals treated with Metformin in comparison to animals treated with vehicle. **F)** Orcein staining shows presence of CCM lesions throughout the CNS in both treatment groups. **G-H)** Analysis in single CCM lesion area and number. **I-J)** analysis in multi-cavernous CCM lesion area and number. N= 7 or 14 mice per group.

### Metformin reverses alterations in the mitochondrial phenotype of brain endothelial cells in chronic *Pdcd10^BECKO^* mice

There is increasing evidence that the chronic nature and exacerbation of CCM disease are partially associated with the activation of CCM endothelial cells by localized neuroinflammation that develops during the CCM pathogenesis and remains unresolved^18,22,23,29,31–35^. This suggests that ongoing vascular inflammation may affect brain endothelial mitochondria during CCM disease, thereby perpetuating the neuroinflammatory response throughout the course of the disease^84^. Moreover, evidence suggests that metformin enhances mitophagy and normalizes mitochondrial function, helping to alleviate inflammation^85^. To understand metformin function, we examined the structure of mitochondria in the brain endothelial cells during CCM disease. We intercrossed the CCM animal model (*Slco1c1-CreERT2;Pdcd10^fl/fl^* or *Slco1c1-CreERT2;Pdcd10^fl/wt^*) with MITO-Tag (B6N.Cg-Gt(ROSA)26Sortm1(CAG-EGFP*)Brsy/J)^86,87^ (Fig. 4A, 4B). This animal model allows conditional expression of an HA-tagged EGFP which localizes to the mitochondrial outer membrane within brain endothelial cells under healthy conditions and during CCM disease (Fig. 4B). Mitochondria in brain vascular endothelial cells were visualized by immunofluorescence using antibodies specific to HA-tagged proteins. The results showed that the mitochondria in brain endothelial cells in the CCM lesions of *Pdcd10^BECKO^*; MITO-Tag mice were smaller and less branched compared to those in the brain endothelial cells of the blood vessels of *Pdcd10^BECKO/wt^*; MITO-Tag control littermates, which do not develop CCM disease (Fig. 4B). To confirm these changes in mitochondria morphology, we performed transmission electron microscopy (TEM) analysis on brain endothelial cells isolated from P60 *Pdcd10^BECKO^* brains and their littermate controls, *Pdcd10^fl/fl^* brains. The TEM analysis showed that mitochondria in the CCM endothelial cells were more rounded, smaller in size, and more numerous than those in the control brain endothelial cells (Fig. 4C, 4D).

**Figure 4.**
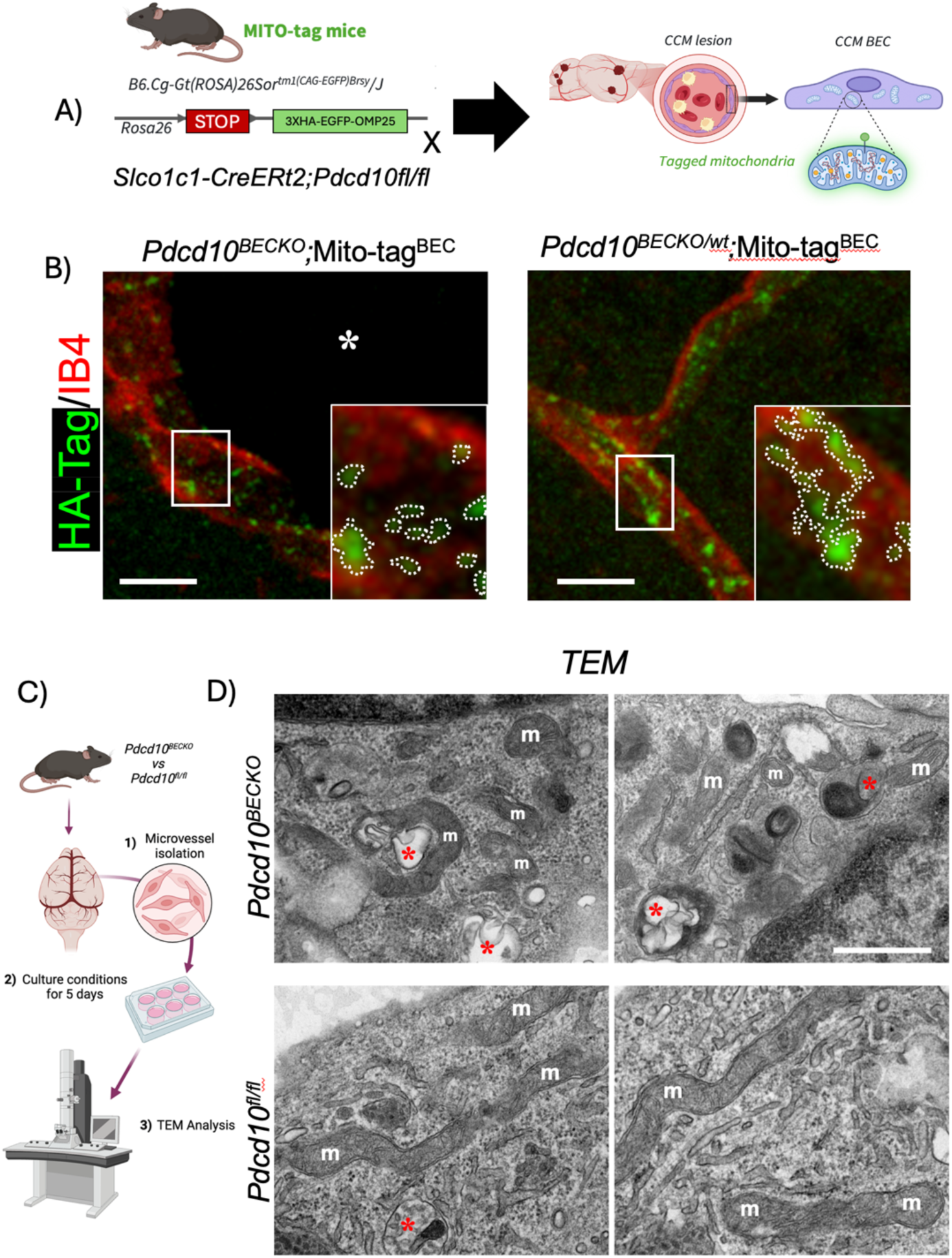
Mitochondria phenotype linked to loss of *Pdcd10* in mouse brain endothelial cells (BECs). A) Animal model used for mitochondria analysis (*Slco1c1-iCreERT2; Pdcd10^fl/fl^; B6.Cg-Gt(ROSA)26Sor^tm1(CAG-EGFP)Brsy^/J*) expresses a hemagglutinin (HA)-tagged enhanced green fluorescence protein (EGFP) that localizes to mitochondrial outer membrane, in a Cre-dependent manner. **B)** Immunofluorescence analysis in a mito-Tag CCM mouse model (*Slco1c1-iCreERT2; Pdcd10^fl/fl^; B6.Cg-Gt(ROSA)26Sor^tm1(CAG-EGFP)Brsy^/J)* showed that mitochondria (HA-Tag, green; white dashed line in the zoomed-in panels) in BECs (Isolectin B4 (IB4), red), are smaller and do no tend to form networks when *Pdcd10* is missing (*Pdcd10^BECKO^*). **C)** Protocol for BEC isolation from Pdcd10BECKO mice and control littermates, cells were maintained in culture conditions for 5 days and then processed for transmission electron microscopy (TEM) analysis. **D)** TEM analysis shows that BEC mitochondria (m) in *Pdcd10^BECKO^* animals are smaller and rounded when compared to BECs in *Pdcd10^fl/fl^* animals. Also, BEC in *Pdcd10^BECKO^* animals seem to present more mitophagy events compared to control animals, as suggested by the proximity of phagosome-like structures to mitochondria (red stars).

Reports suggest that Metformin activates AMPK, enhances mitophagy, and normalizes mitochondrial function^85,88^. AMPK is vital for mitochondrial fission under energy stress, and its pharmacological activation can eliminate damaged mitochondria, maintaining cellular homeostasis^89^. However, the roles of these mechanisms and the impact on inflammatory signaling in brain vascular malformations remain unexplored. Next, we next assessed the effects of metformin on mitochondrial morphology in brain endothelial cells isolated from chronic *Pdcd10^BECKO^*;MITO-Tag mice, as these cells displayed altered mitochondrial phenotypes (Fig. 4). We isolated brain endothelial cells from *Pdcd10^BECKO^*;MITO-Tag mice and *Pdcd10^wt/wt;^* MITO-Tag littermates and incubated them with metformin (50 µM) or vehicle for 72 hrs, followed by fixation and post-analysis to assess mitochondrial structure (Fig. 5A). Brain endothelial cells from *Pdcd10^BECKO^;* MITO-Tag mice treated with metformin exhibited a reversal in several mitochondrial phenotype parameters when compared to vehicle-treated brain endothelial cells from the same *Pdcd10^BECKO^*;MITO-Tag mice. The treatment resulted in an increase in both the area and perimeter of the mitochondria, as well as a more elongated mitochondrial shape (Fig. 5B-D, Supplemental Fig. 2). Additionally, the complexity of the mitochondrial networks increased under treatment with metformin, with values comparable to those observed in brain endothelial cells from control mice (Fig. 5F-5H). Excessive mitochondrial fragmentation in endothelial cells has been linked to vascular inflammation^90^. We observed that brain endothelial cells isolated from chronic *Pdcd10^BECKO^*;MITO-Tag mice that had mitochondrial fragmentation also exhibited increased levels of VCAM-1, which is a marker of inflammation (Fig. 5J, 5K). Furthermore, treatment with metformin resulted in a significant reduction of VCAM-1 expression in brain endothelial cells of *Pdcd10^BECKO^*;MITO-Tag mice (Fig. 5K). These results indicate that metformin treatment directly affects brain endothelial cells by reversing mitochondrial phenotype and reducing VCAM-1 expression associated with CCM disease. This could represent an important strategy to decrease the CCM lesion burden seen in animal models.

**Figure 5.**
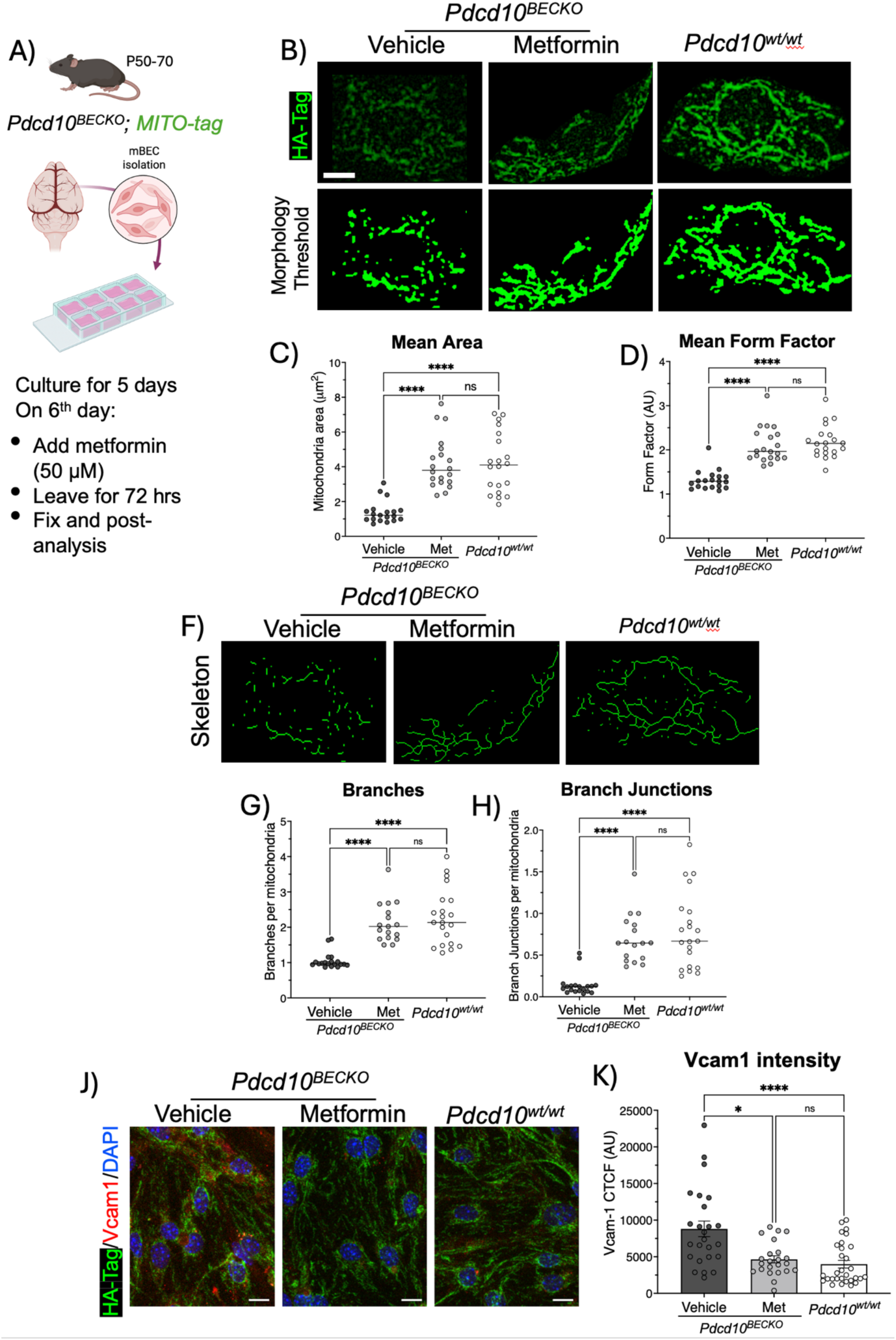
Metformin reverses changes in mitochondria structure and endothelial inflammation caused by loss of brain endothelial *Pdcd10*. **A)** Schematic diagram illustrating the protocol that utilizes the BECs expression MITO-tag from a CCM animal model. **B)** BECs isolated from *Pdcd10^BECKO^* mice were treated with Metformin or vehicle and stained with antibodies against HA-tag to label mitochondria (green). The threshold set up defines the elements (mitochondria) to be analyzed based on the antibody staining to assess mitochondria shape. **C)** The mean mitochondrial area and **D)** the mean form factor indicate the presence of smaller, more rounded, and fragmented mitochondria in brain endothelial cells from *Pdcd10^BECKO^* animals. Treatment with metformin reversed these mitochondrial changes. **F)** Skeletons of the mitochondria networks categorized by the same threshold showed in *A.* The analysis of **G)** branch and **H)** branch joints indicates that the complexity of the mitochondria network is reduced in BECs isolated from *Pdcd10^BECKO^* mice. However, treatment with metformin reversed the defects observed in the mitochondria network, bringing it back to a level similar to that found in control BECs from *Pdcd10^wt/wt^* mice. **J, K)** BECs from *Pdcd10^BECKO^* animals showed a significant increase in VCAM1 expression when compared to control BECs from *Pdcd10^wt/wt^* mice. However, VCAM1 intensity was significantly reduced to control levels when cells from *Pdcd10^BECKO^* animals were treated with metformin. Data are mean +/- SEM, N= 20 cells from 3 biological replicates.

### AMPK activator PF-06409577, reverses alterations in the mitochondrial phenotype of brain endothelial cells in chronic *Pdcd10^BECKO^* mice

We hypothesize that metformin reverses the brain endothelial mitochondrial phenotype associated with CCM disease through the activation of AMPK. Therefore, we next tested whether an AMPK activator is sufficient to reverse the mitochondrial phenotype in brain endothelial cells derived from chronic *Pdcd10^BECKO^*;MITO-Tag mice (Fig. 4, Fig. 5). Brain endothelial cells were isolated from *Pdcd10^BECKO^*;MITO-Tag mice and *Pdcd10^wt/wt^*;MITO-Tag littermates, and incubated with either PF-06408677 (0.1 µM) or vehicle. After 72 hours, we performed fixation and post-analysis to assess mitochondrial structure (Fig. 6A-6F, Supplemental Fig. 3). In line with the results observed with metformin, we found that brain endothelial cells from *Pdcd10^BECKO^*;MITO-Tag mice treated with PF-06408677 showed a significant improvement in several mitochondrial phenotype parameters compared to vehicle-treated brain endothelial cells from the same *Pdcd10^BECKO^*;MITO-Tag mice. Moreover, treatment with PF-06408677 led to a significant reduction of VCAM-1 expression in the brain endothelial cells of *Pdcd10^BECKO^*;MITO-Tag mice, indicating that reversing the mitochondrial phenotype strongly alleviates VCAM-1 expression (Fig. 6G, 6H). In summary, our study reveals that AI can be utilized as a tool to identify novel therapeutic targets aimed at enhancing endothelial adaptation or reversing disease phenotypes with vascular malformations. Moreover, metformin, an AMPK agonist and mTOR inhibitor, can reverse the changes in cell-cell junction organization and the increased levels of KLF4 protein found in human CCM endothelial in vitro model. Results also revealed that brain endothelial cells in chronic CCM mouse models exhibit altered mitochondrial phenotypes associated with elevated levels of the inflammatory marker VCAM-1. Furthermore, we found that both metformin and a potent AMPK activator, PF-06409577, can reverse these mitochondrial changes in brain endothelial cells and reduce the elevated VCAM-1 levels linked to chronic neuroinflammation in CCM disease. We propose that metformin may offer endothelial cytoprotection and enhances metabolic adaptation in response to brain vascular malformations associated with neuroinflammation through the activation of AMPK (Fig. 6I).

**Figure 6.**
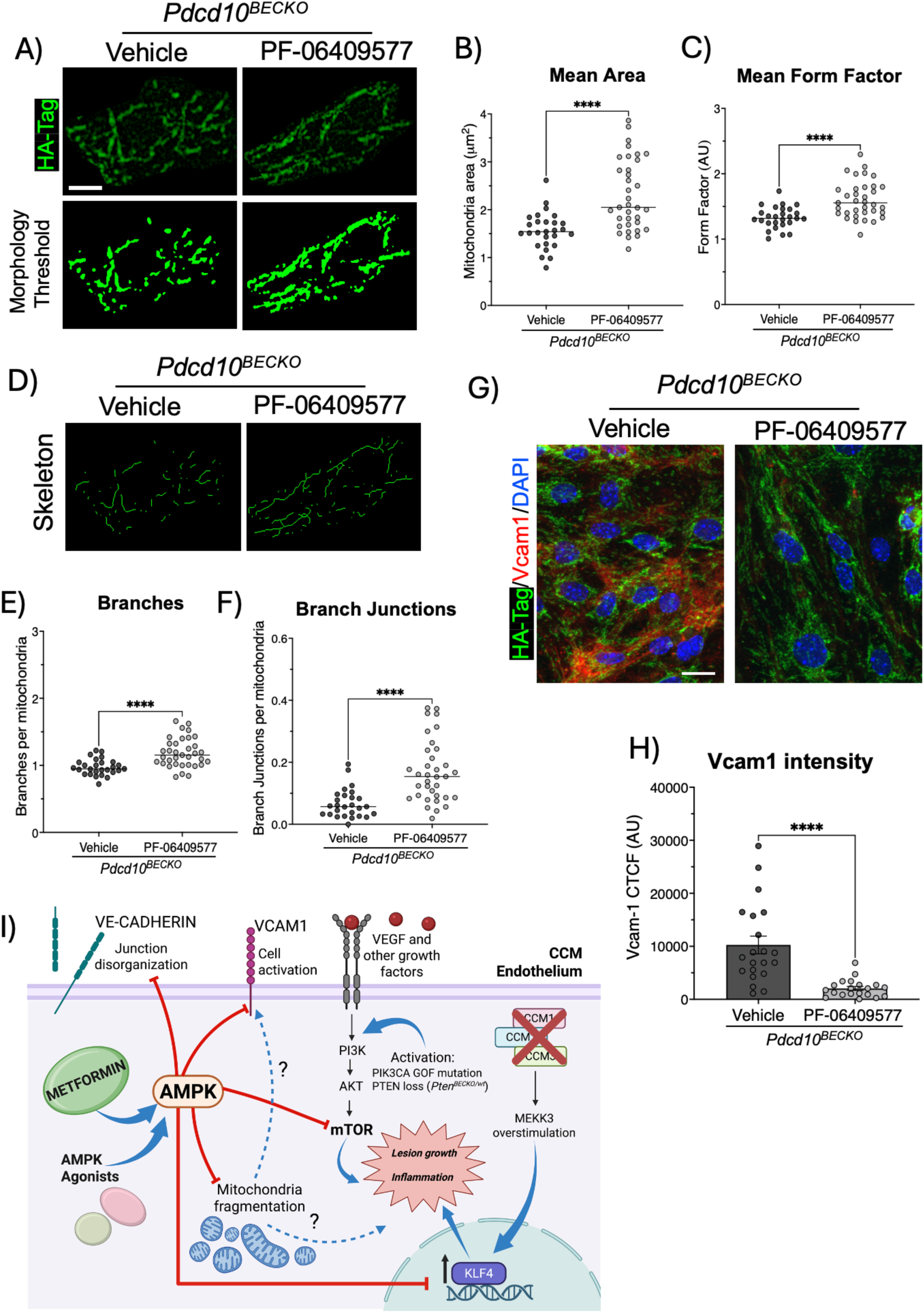
AMPK agonist, PF-06409577, reverses changes in mitochondria structure and endothelial inflammation caused by loss of brain endothelial *Pdcd10*. **A)** Brain endothelial cells isolated from *Pdcd10^BECKO^* mice were treated with PF-06409577 or vehicle and stained with antibodies against HA-tag to label mitochondria (green). The threshold set up defines the elements (mitochondria) to be analyzed based on the antibody staining to assess mitochondria shape. **B)** The mean mitochondrial area and **C)** the mean form factor indicates that brain endothelial cells from *Pdcd10^BECKO^* treated with an AMPK agonist reversed mitochondrial changes associated with smaller, rounded, and fragmented mitochondria observed in *Pdcd10^BECKO^*. **D)** Skeletons of the mitochondria networks categorized by the same threshold showed in *A.* The analysis of **E)** branch and **F)** branch joints indicates that the complexity of the mitochondria network is reduced in brain endothelial cells isolated from *Pdcd10^BECKO^* mice. However, treatment with PF-06409577 reversed the defects observed in the mitochondria network. **G, H)** BECs from *Pdcd10^BECKO^* animals showed high VCAM1 expression, which was significantly reduced when cells were treated with the AMPK agonist. **I)** Schematic representation of the proposed mediation of metformin and AMPK agonists (i.e. PF-06409577) in rescuing of junction disorganization, cell activation, mitochondria fragmentation and mTOR overactivation through AMPK activation in CCM endothelial cells. Data are mean +/- SEM, N= 20 cells from 3 biological replicates.

## Discussion

Here we used artificial intelligence (AI), high-content screening, pharmacokinetics, pharmacodynamics, and toxicokinetic to identify potential long-term drug treatments for CCM. The validation of AI and high-content screening highlighted effective strategies for pinpointing new therapeutic targets in CCM pathology, as shown by improving metabolic adaptation in response to signaling pathways activated during endothelial vascular malformations such as hypoxia and inflammation^18,22,23,35^. This study showed that the FDA-approved drug metformin, an AMPK agonist and mTOR inhibitor, may function as a potential treatment for CCM. Metformin treatment reduces neurovascular lesions in both aggressive and chronic CCM mouse models.

The reduction in lesion burden was associated with an enhanced AMPK activation and decreased mTOR signaling. These effects were evidenced by increased levels of AMPK phosphorylation and decreased S6 ribosomal protein phosphorylation, which served as pharmacodynamic markers. Pharmacokinetic and toxicological studies in a chronic CCM animal model indicated that long-term administration of a therapeutic dose of metformin results in a safe profile. Our study demonstrated that brain endothelial cells in chronic CCM mouse models exhibit significant mitochondrial fragmentation and increased levels of VCAM-1 expression. We also discovered that metformin, as well as a potent AMPK activator, PF-06409577, can reverse the mitochondrial phenotype in brain endothelial cells and reduce the elevated VCAM-1 levels associated with chronic CCM disease. These findings suggest that metformin may offer endothelial cytoprotection and reverse the CCM endothelial phenotype through an increased in AMPK activation and reversal of mitochondrial alterations. Thus, metformin may enhance metabolic adaptation to brain vascular malformations by activating AMPK.

CCMs are neurovascular lesions that affect both adults and children, whose morbidity often results from thrombosis, bleeding, and endothelial dysfunction, which can have debilitating effects on patients^4,23,53,91,92^. Notably, the various types of CCMs (CCM1, *CCM2*, or *CCM3*^1,5,93,94^), whether sporadic or familial, are generally considered histologically indistinguishable^15^. This suggests that they may have either similar or distinct pathological mechanisms, with a common outcome that entails the transition from individual lesions to active late-stage multi-cavernous lesions, which are closely mirrored in chronic CCM mouse models^23,35,82,95–97^. Currently, there are no effective therapeutic options available for preventing or treating CCMs, making surgery the only viable option^5,54,57^. However, surgery may not be feasible for lesions located in critical or hard-to-reach areas or in cases where patients have multiple lesions throughout the brain and spinal cord. As a result, there is a significant need for long-term therapies for CCMs, as existing medications only provide partial relief for some symptoms^5–10^. Notably, 60-70% of surgically resected CCM1 have acquired a somatic, activating mutation in PI3KCA or MAP3K3^7–9,55,56^. Mouse models have shown that CCM1 aggressiveness can be caused by gain-of-function mutations in *PIK3CA* in endothelial cells, and that the mTOR inhibitor rapamycin can prevent CCM formation^9,79^. However, the doses of rapamycin (100 or 400 ug/day) used in that preclinical study^9^ would be expected to achieve supratherapeutic blood levels that would not be safely tolerated in humans (as doses produce blood concentrations >15 ng/mL) and may cause serious side effects^98^. Therefore, additional studies are required to determine the optimal oral dosing of rapamycin^98^, minimize the raised safety concerns regarding long-term use as a potential immunosuppressor, or identify other better-tolerated mTOR inhibitors.

Our study reveals that AI can be used to decipher complex biological programs underlying vascular malformations and to identify novel therapeutic targets. This was achieved by integrating extensive and diverse data from various biomedical sources^60^ with CCM transcriptomic studies^15,21,23,35,68^. In this study, we included therapeutic targets that have acceptable safety profiles and FDA-approved drugs suitable for potential repurposing in the long-term treatment of CCM disease. We compiled a list of 71 drugs for experimental validation, several of which specifically target cell metabolism, inflammation, cell growth, proliferation, mitochondrial functions, and fatty acid regulation, among others. While we use a wide range of doses to test pharmacological compounds, some drugs may require further dose optimization to confirm biological effects on CCM disease. We focused on metformin because it has a well-documented safety profile for long-term use in both children and adults^70,71^. Additionally, it can help reduce mTOR signaling^99^ by activating AMPK^100–102^, which we also observed in pre-clinical CCM lesions in animal models. Metformin also has excellent properties, including high solubility, permeability, stability, and favorable pharmacokinetic and pharmacodynamic profiles. However, the limited lack of information regarding metformin concentration and safety profiles for brain-related conditions such as CCMs. Here, we evaluated pharmacologically relevant dosing regimens, preclinical efficacy, and safety of metformin in treating CCM disease. Although we noted efficactiveness and did not observe toxicity when using metformin (100 mg/kg/day) for long-term treatment in adult CCM3 animal model, further studies on dose effects and treatment protocol are necessary to effectively translate preclinical findings into human phase trials of metformin therapy for CCM patients. Furthermore, it is important to note that individuals with severe CCM1 disease, particularly those with PIK3CA gain-of-function mutations^7–9^, will likely require higher treatment doses than those with CCM1 alone. This is attributed to an early and excessive increase in PI3K-mTOR signaling^9,23^. It would also be beneficial to investigate the metformin dosage requirements in chronic models across CCM1 and CCM3. Additionally, exploring combinations of metformin with other robust pharmacological treatments, such as rapamycin, could be valuable. For example, rapamycin could be administered for a short term, followed by long-term metformin therapy. Understanding different therapeutic approaches will help develop personalized metformin treatments for individuals with CCM disease.

Metformin, the most widely prescribed oral anti-diabetic medication, has various effects on different signaling pathways^73,103^. Some of its effects are unrelated to glycemic control, such as inhibiting PI3K-AKT-mTOR signaling^99,104,105^ and preventing activation of NLRP3 inflammasome^106–108^ in cancers and during inflammation. It also prevents cardiovascular diseases through both AMPK-dependent and AMPK-independent mechanisms^109^. Reports suggest that Metformin activates AMPK, enhances mitophagy, and normalizes mitochondrial function^85,88^. Notably, maintaining a healthy mitochondria population is crucial for the proper functioning of the brain endothelial cells^110–112^. We observed that CCM endothelial cells exhibit an altered mitochondrial phenotype, which was associated with elevated VCAM1 levels as an inflammatory marker in CCMs^23^. This could contribute to the persistent inflammation in brain endothelial cells observed in CCM disease^18^ by releasing mitochondrial components and metabolic products that could trigger inflammatory responses when they enter the cytosol or the extracellular environment^84^. Future studies should investigate whether metformin affects mitochondrial autophagy, also known as mitophagy. This process involves dysfunctional mitochondria being enclosed by autophagosomes and transported to lysosomes for degradation. Mitophagy enables cells to repair themselves and respond to stress, ultimately preventing unnecessary cell death^113,114^. This process is vital for brain endothelial cells, which are primarily post-mitotic and cannot be easily replaced. Additionally, AMPK activation is crucial for mitochondrial fission during energy stress. Its pharmacological activation can eliminate damaged mitochondria, thereby helping maintain cellular homeostasis^89^. Therefore, studying how AMPK contributes to metabolic adaptation in vascular malformations through these mechanisms is an important area for future research. It is also important to consider that metformin is a pleiotropic molecule that may influence endothelial cells and immune cells^74,85,107,115^. While our main focus was on how metformin affects brain endothelial cells in mouse models of CCM disease, we cannot exclude the possibility that metformin may also influence peripheral immune cells, such as macrophages, neutrophils, and lymphoid cells^23,31^. These inflammatory immune cells exhibit increased mTOR activity (data not shown) and play a role in the development of CCM disease^23,31–33^. Our study used AI analysis, high-content screening, toxicology assessments, and pharmacokinetic studies in CCM models to evaluate the safety, efficacy, toxicity, and mechanisms of metformin’s action in CCM disease. We propose that metformin activates AMPK, which enhances metabolic adaptation to vascular malformations by inhibiting mTOR and reversing brain endothelial mitochondrial phenotypes associated with inflammatory activity. Metformin has potential as a pharmacological treatment for long-term therapy of CCM disease, which could have significant therapeutic implications for affected individuals.

## Authors’ contributions

E.F-A. H.G-G., C.B., designed and performed the experiments, analyzed and interpreted the data, generated the figures, and wrote the manuscript. E.F- A. designed and prepared all illustrations. I.R.N performed the TEM analysis. B.N., A.S., Z.M., N.C., helped with animal experiments, genotypes and interpretation of results. I.A.A. J.S., B.G., H.H.P., J.T., J.D. M., provided critical reagents and interpretation of results and helped to review the manuscript. A.T., B.K., M.A.L-R. Contributed to the AI-CCM analysis and triage. M.A.L-R. conceived the project, designed the overall study, analyzed and interpreted data, and wrote the manuscript.

## Acknowledgments

The Benevolent target ID product and technology team James Dunbar, Gabi Griffin, Amparo Toboso-Navasa, Chun Wan, Chris Tame, Dennis Schwartz, Mara Tatari, Kirill Shkura. The authors thank Mark H. Ginsberg, Angioma Alliance including Amy Akers, Connie Lee and Angela Glading for helpful discussion.

---

**Supplemental Table 1.**
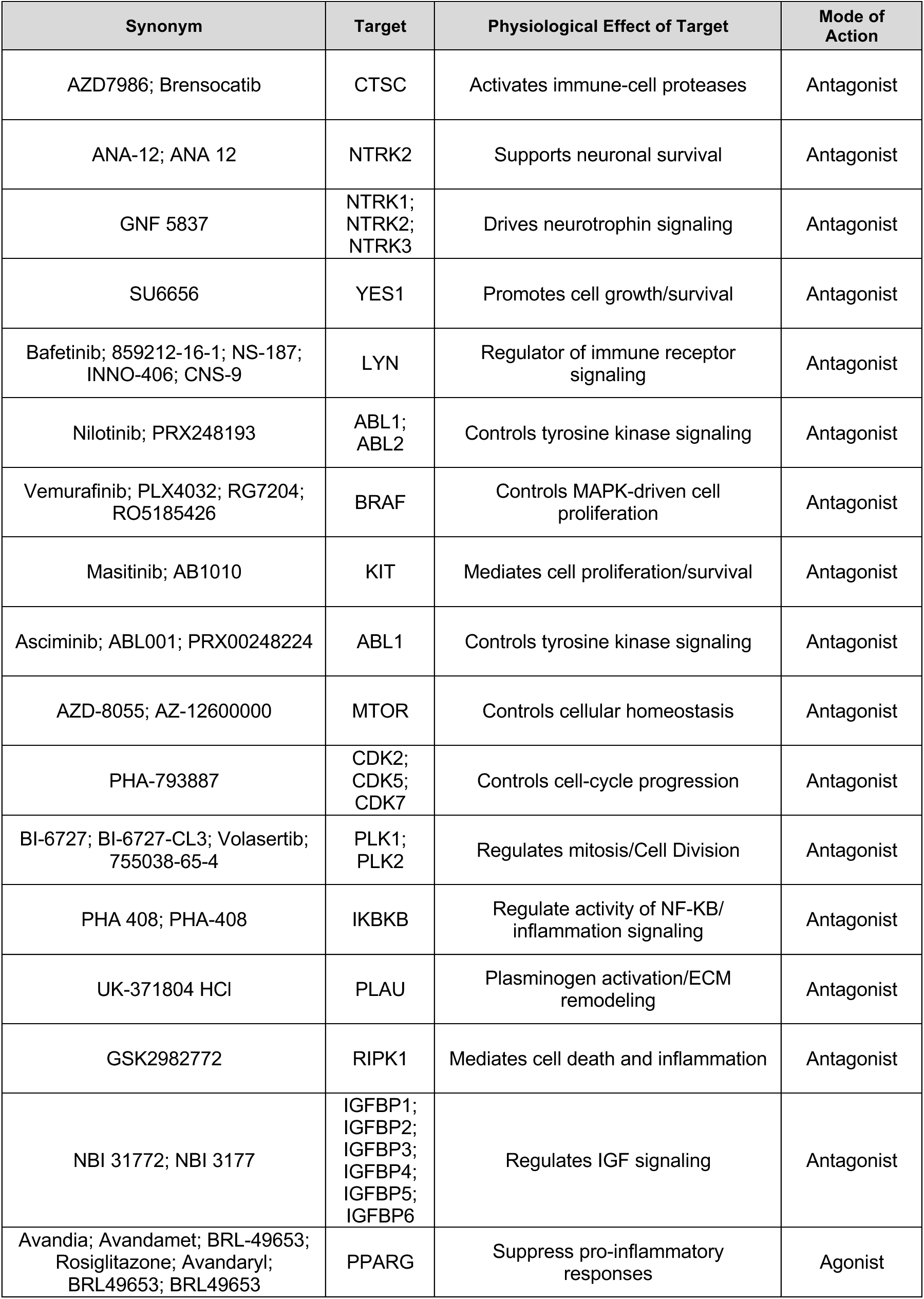
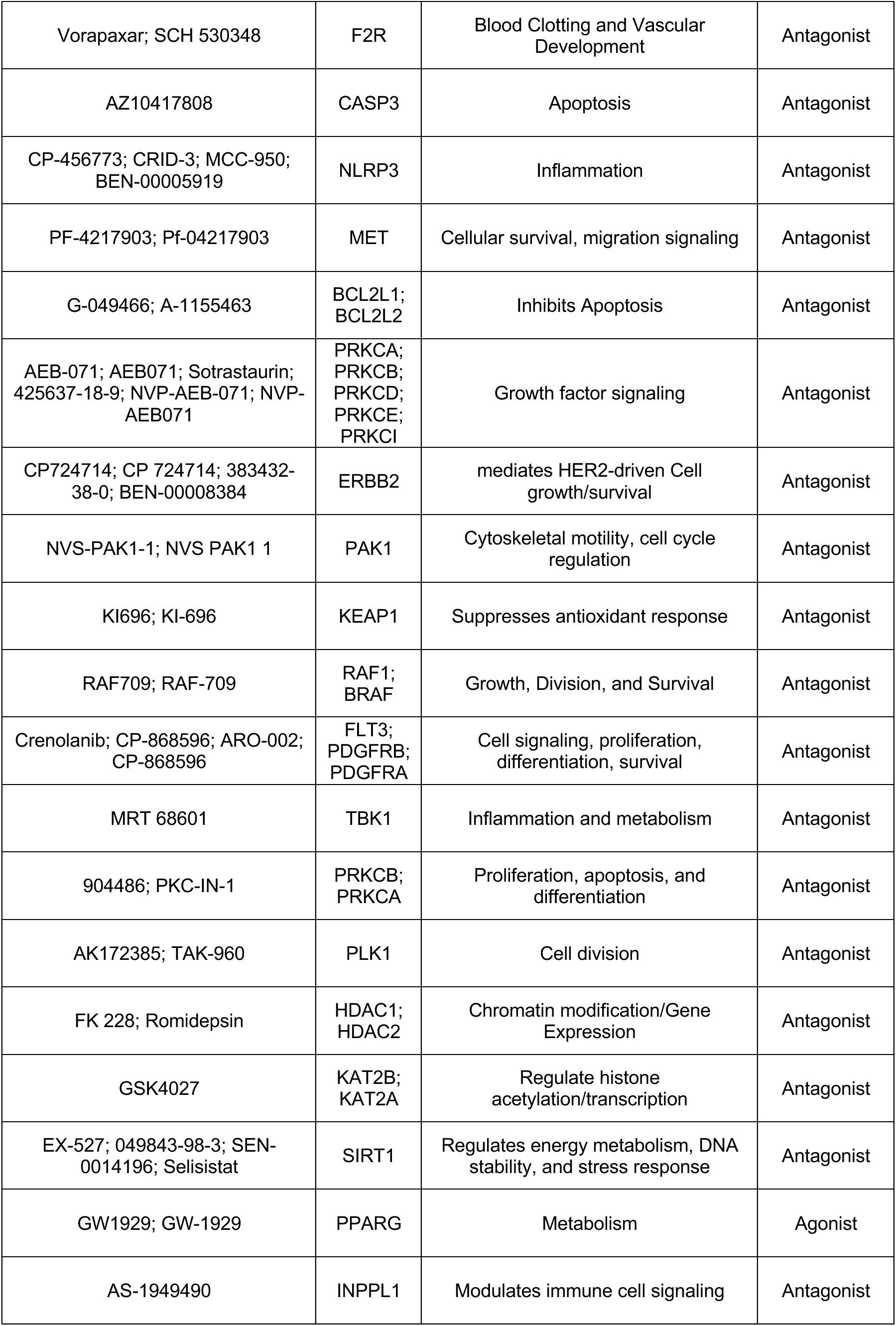
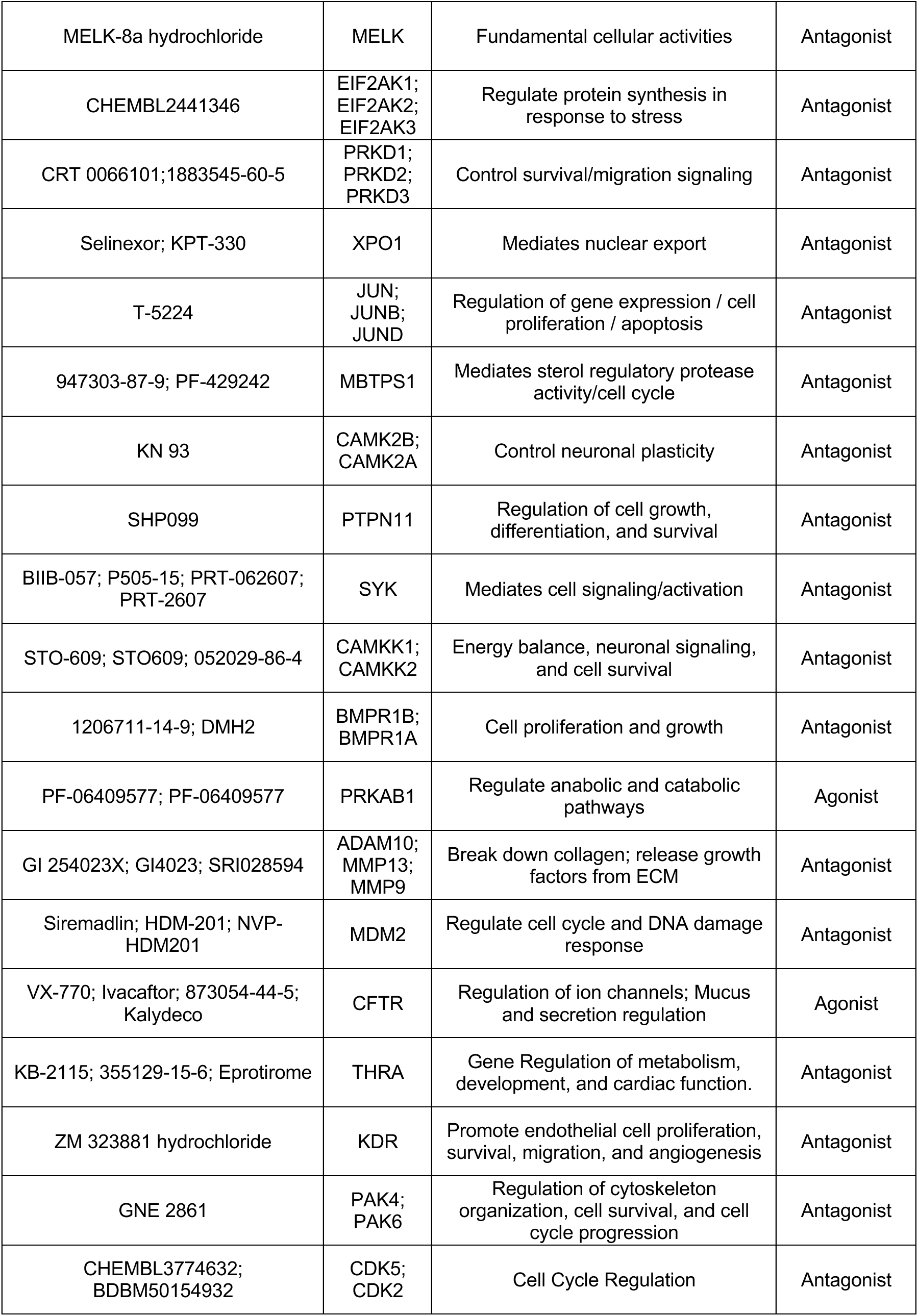
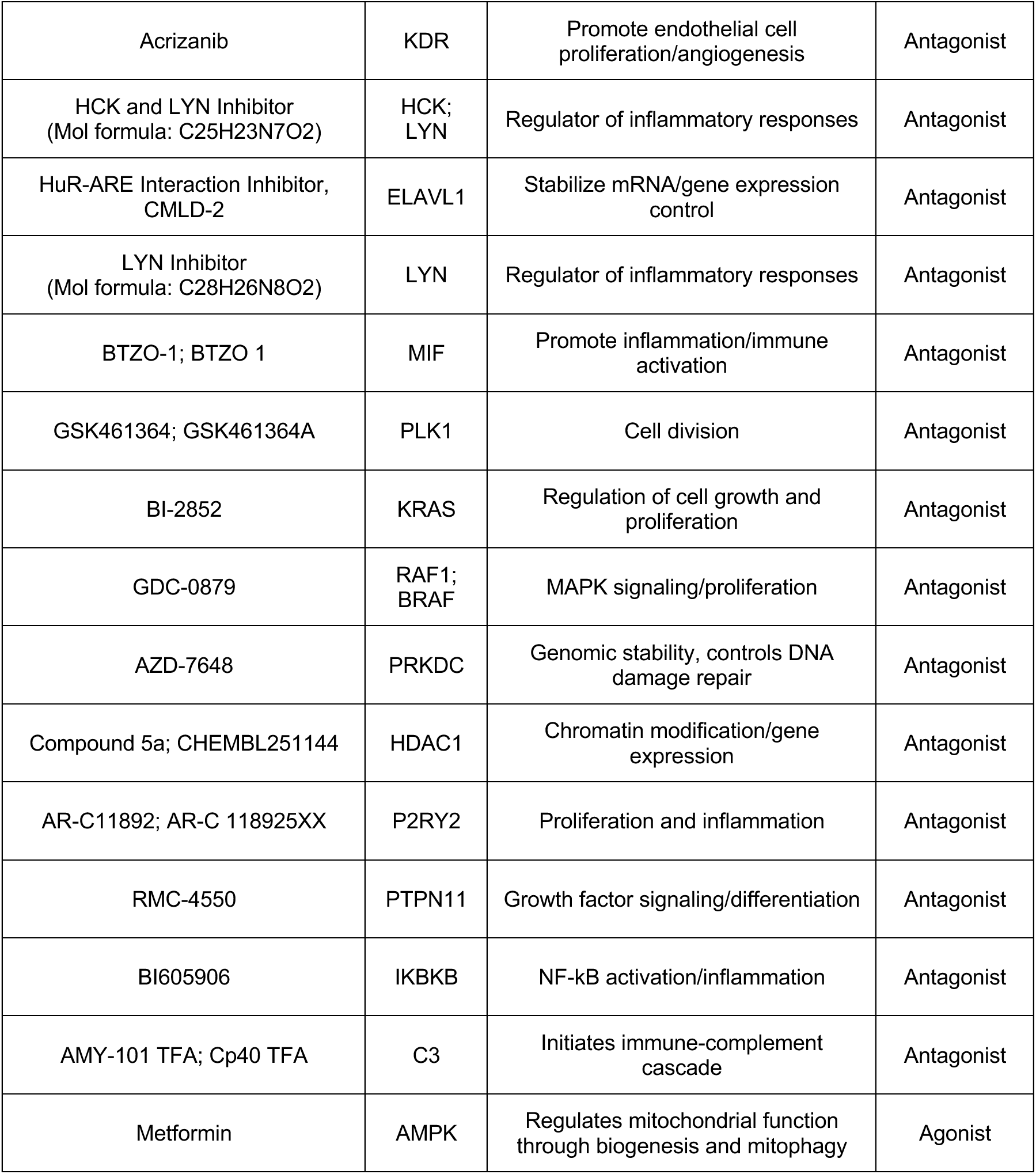
Drug screening compounds and their targets for CCM treatment approach.

**Supplemental Figure 1.**
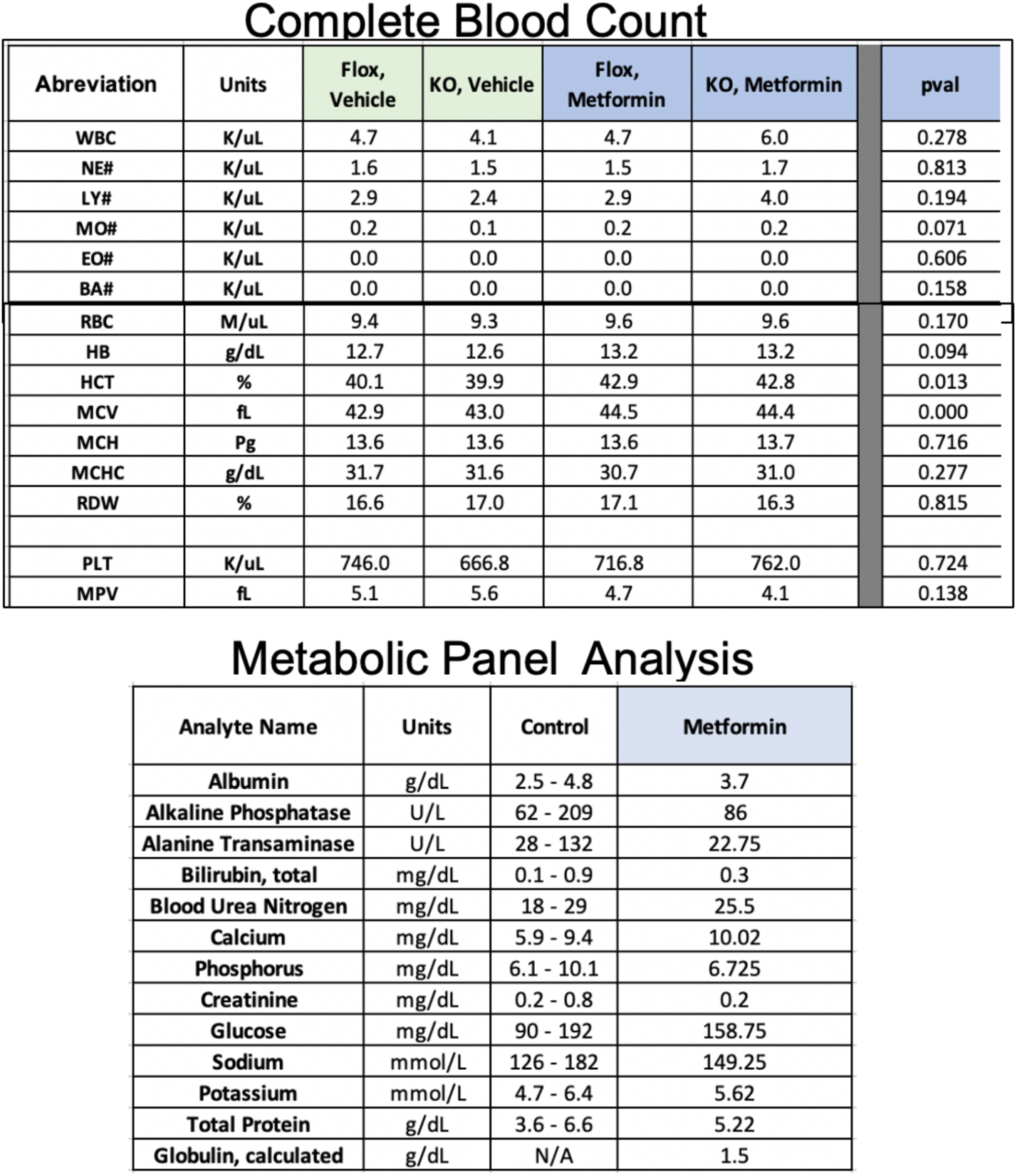
Toxicological assessment. Complete Blood Count and Metabolic Panel Analysis in *Pdcd10^BECKO^*(KO, CCM3) and littermate controls *Pdcd10^flox^*^(^Flox) following treatment with Metformin (100 mg/Kg/day) for 20 days (n=3 or 4).

**Supplemental Figure 2.**
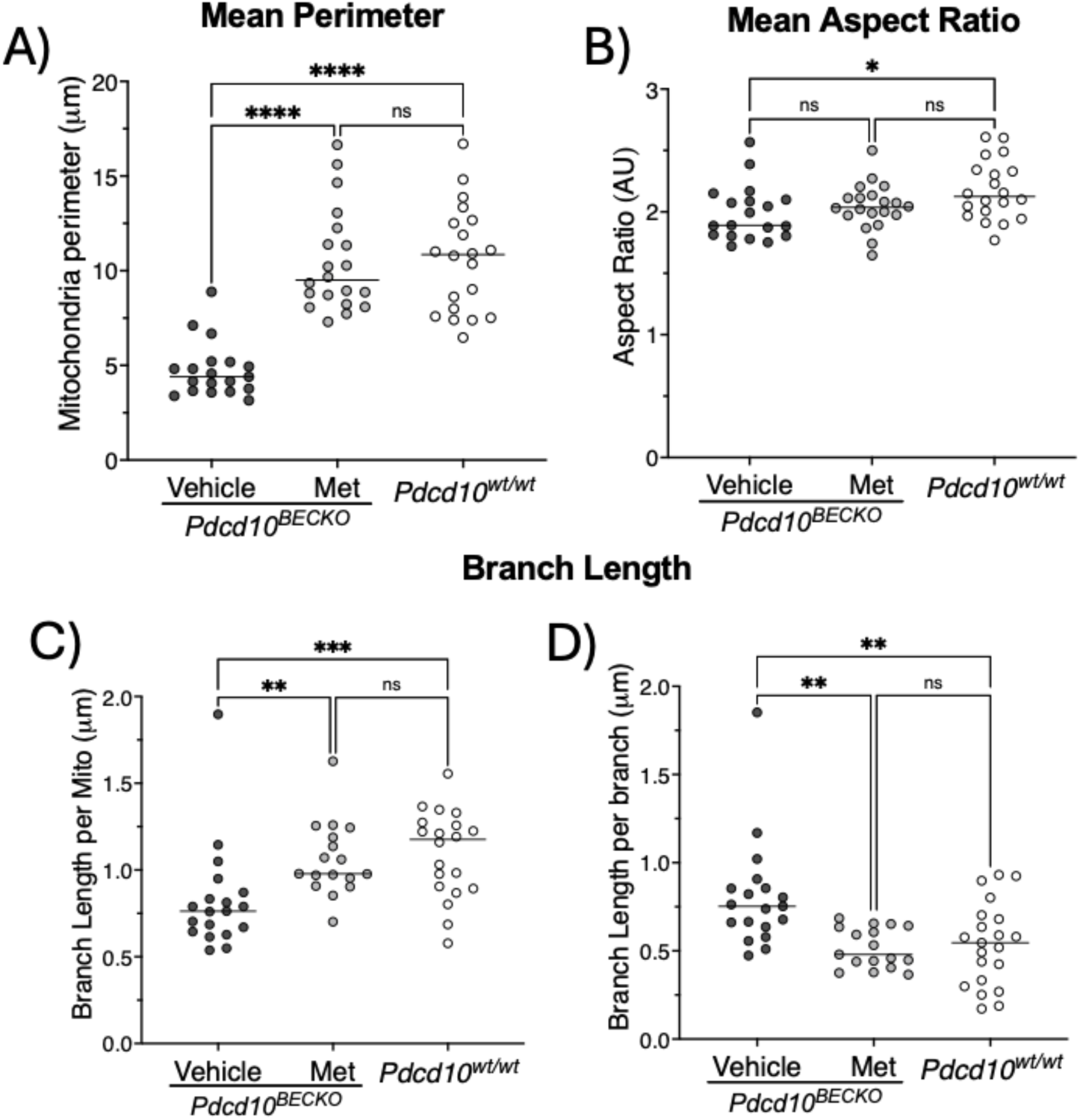
Metformin reverses changes in mitochondria perimeter, aspect ratio and network complexity caused by loss of brain endothelial *Pdcd10*. BECs isolated from *Pdcd10^BECKO^* mice were treated with Metformin or vehicle. **A)** The mean mitochondrial perimeter and **B)** the mean aspect ratio (elongation parameter) indicates that BECs from *Pdcd10^BECKO^* are smaller, more rounded, and show appearance of fragmented mitochondria. In addition, the analysis of **C)** branch length per mitochondria and **D)** branch length per branch indicate that the complexity of the mitochondria network is significantly reduced in BECs isolated from *Pdcd10^BECKO^* mice when compared to BECs isolated from control littermates. Treatment with metformin was shown to reverse the changes in the mitochondria network linked to loss of *Pdcd10*, bringing it back to a level similar to that found in control BECs from *Pdcd10^wt/wt^* mice. Data are mean +/- SEM, N= 20 cells from 3 biological replicates.

**Supplemental Figure 3.**
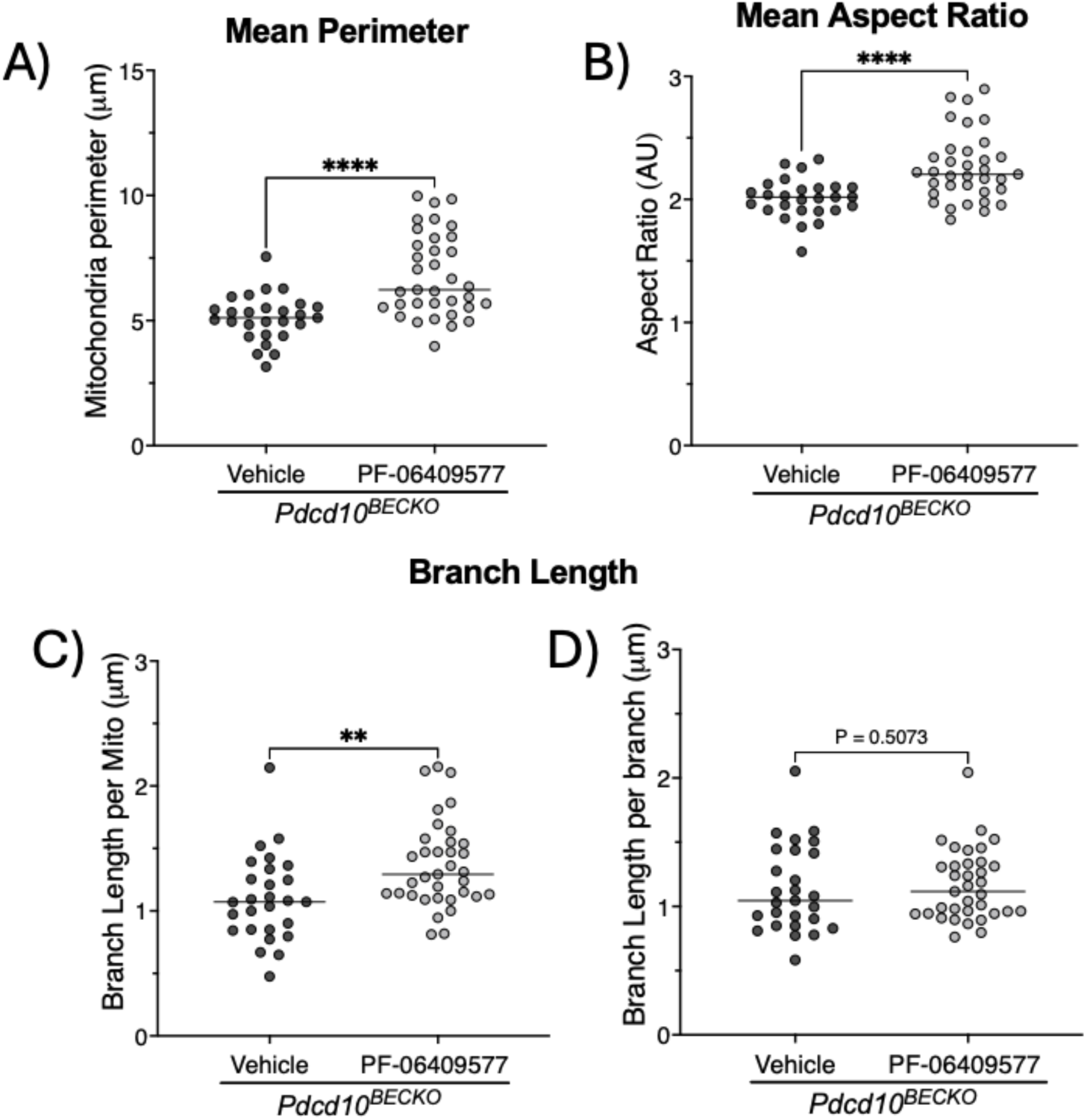
AMPK agonist, PF-06409577, reverses changes in mitochondria perimeter, aspect ratio and network complexity caused by loss of brain endothelial *Pdcd10*. BECs isolated from *Pdcd10^BECKO^* mice were treated with PF- 06409577 or vehicle. **A)** The mean mitochondrial perimeter and **B)** the mean aspect ratio indicates that brain endothelial cells from *Pdcd10^BECKO^* treated with PF-06409577 reversed mitochondrial changes associated with smaller, rounded, and fragmented mitochondria observed in BECs from *Pdcd10^BECKO^*. The analysis of **C)** branch length per mitochondria and **D)** branch length per branch indicate that the mitochondrial network complexity is reduced in brain endothelial cells isolated from *Pdcd10^BECKO^* mice. However, treatment with PF-06409577 reversed the defects observed in the mitochondria network. Data are mean +/- SEM, N= 20 cells from 3 biological replicates.

